# A guide to computational cotranscriptional folding featuring the SRP RNA

**DOI:** 10.1101/2023.06.01.543211

**Authors:** Stefan Badelt, Ronny Lorenz

## Abstract

Although RNA molecules are synthesized via transcription, little is known about the general impact of cotranscriptional folding in vivo. We present different computational approaches for the simulation of changing structure ensembles during transcription, including interpretations with respect to experimental data from literature. Specifically, we analyze different mutations of the E.coli SRP RNA, which has been studied comparatively well in previous literature, yet the details of which specific metastable structures form, as well as when they form are still under debate. Here, we combine thermodynamic and kinetic, deterministic and stochastic models with automated and visual inspection of those systems to derive the most likely scenario of which substructures form at which point during transcription. The simulations do not only provide explanations for present experimental observations, but also suggest previously unnoticed conformations that may be verified through future experimental studies.

## 1 Introduction

During RNA transcription, a polymerase adds single nucleotides to the 3’ end of the molecule. The rate at which nucleotides are attached varies from 20 to 200 nt s^−1^ (nucleotides per second), but the processes is not necessarily continuous: pausing during transcription can last for multiple seconds [1].

In comparison, the fastest formations of helices can be on the order of microseconds, so-called branch migration processes happen on the order of milliseconds, and large structural rearrangements may take seconds, hours, or much longer. As a consequence, the 5’ end of RNA molecules can form secondary structures temporary during transcription, and those structures may either remain stable for some time, or fold into another (potentially thermodynamically optimal) secondary structure.

Presumably, cotranscriptional folding can be used as a mechanism to encode multiple functions into a single molecule, or simply to encode a delay before an RNA molecule switches into its active conformation. However, to which extent nature harnesses the potential of cotranscriptional folding is still largely unknown. It is possible that many RNA molecules must form exactly one thermodynamically stable structure, and that sequences (as well as interactions with other cellular components) evolve to minimize the effects of cotranscriptional folding traps.

In the last decades, the potential importance of cotranscriptional folding has been confirmed experimentally e.g. the work reviewed by Schärfen and Neugebauer [2], but it is still remarkably difficult to pin down the relevant intermediate structures that a given molecule exhibits during transcription.

This chapter demonstrates how RNA folding software can be used to investigate the effects of cotranscriptional folding, and assist with the interpretation of experimental data. We describe a computational analysis of cotranscriptional folding at different levels of detail using existing RNA folding software. Specifically, we compare minimum free energy (MFE) folding, stochastic sampling from the equilibrium distribution, stochastic sampling from the cotranscriptional distribution and a deterministic model of cotranscriptional folding.

## 2 Material

### 2.1 Hardware

To follow our tutorial, command line programs have to be executed on a computer providing a Unix terminal. We recommend using a `Linux` or `Mac` operating system, but also the `Windows subsystem for Linux (WSL)` is possible (see Note 1).

### 2.2 Software

#### 2.2.1 Scientific software

The required software is free for educational and scientific use and the source code can be obtained from the following resources:

1. `ViennaRNA Package` [3] http://www.github.com/ViennaRNA/ViennaRNA
2. `Kinfold` [4] http://www.tbi.univie.ac.at/RNA/Kinfold
3. `DrTransformer` [5] http://www.github.com/ViennaRNA/drtransformer
4. `DrForna` [6] http://viennarna.github.io/drforna/

For a smooth installation experience (especially regarding `ViennaRNA Package Python` bindings), we recommend using `bioconda` [7], rather than compilation from source. We show step-by-step instructions in Sec. 3.2. The API reference manual for the `ViennaRNA Package` is available at https://www.tbi.univie.ac.at/RNA/ViennaRNA/doc/html/index.html. It provides a more technical description of individual functions, especially regarding the `ViennaRNA Package Python` bindings used throughout this tutorial.

#### 2.2.2 Dependencies

The following scripting languages and packages are used in this tutorial:

1. `Python>=3.8` – https://www.python.org/
  a. `pandas` for data processing [8]
  b. `matplotlib` for plotting data [9]
2. `R` – https://www.r-project.org/ [10]
  a. `optparse` for command line argument parsing
  b. `ggplot2` for plotting the data [11]
  c. `viridis` for color-blind friendly palettes [12]
  d. `reshape2` and `scales` for various internal data formatting [13]
3. `git` (optional) – https://git-scm.com/
4. `conda` (optional) – https://conda.io/

### 2.3 Data and scripts

The files provided for following this tutorial are:

1. four different SRP sequences [14, 15],
2. cotranscriptional SHAPE reactivity data files [16, 15], and
3. scripts for data processing, file format conversion, analysis and plotting.

We provide all required files and scripts in a `git` repository at https://github.com/ViennaRNA/DrTutorial.

## 3 Methods

### 3.1 Notation

The instructions for execution of command line programs or scripts in a `Unix` terminal are wrapped in boxes and always preceded with “>“, a greater sign that is *not* part of the program call:

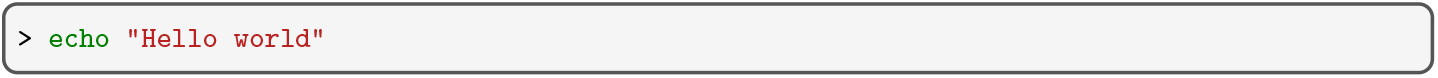

`Python` code is presented in boxes with line numbers, e.g.:

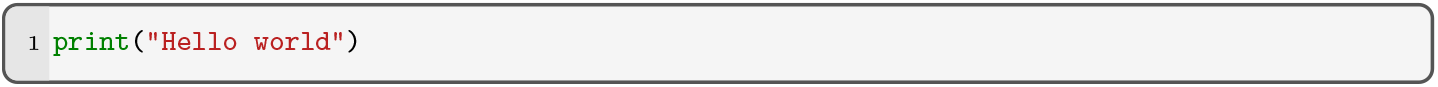

### 3.2 Installation

Instead of compiling and installing the required software in Sec. 2.2 from source, the ready-made binary packages can be directly installed via `conda`. After `conda` is installed and activated, e.g. see https://conda.io/, the following channels must be set – in this order – to obtain the latest versions of the software:

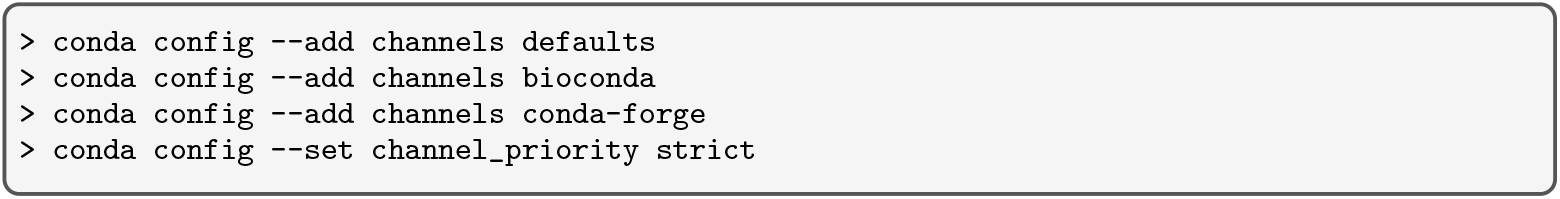

The `ViennaRNA Package` (including `Kinfold`) and `DrTransformer` are installed using the following commands:

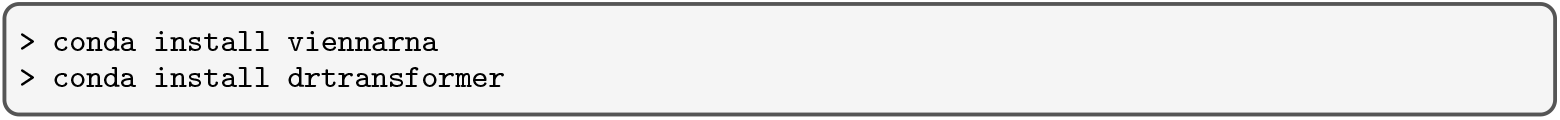

### 3.3 Setup of the data and script repository

All data files and scripts used throughout this chapter are provided in a git repository available at https://github.com/ViennaRNA/DrTutorial. We recommend using `git` to get a local copy of these files, i.e. by cloning the repository:

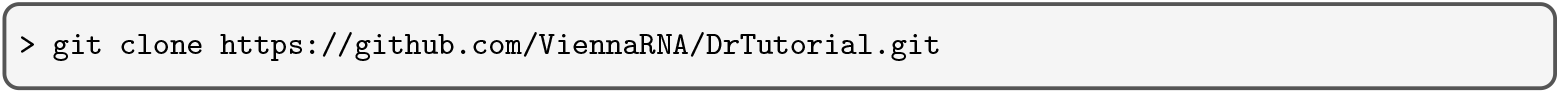

Using `git` has the main advantage that users can contribute to future versions of this tutorial. Alternatively, the directory can be downloaded and extracted from the corresponding `ZIP` archive file. Both ways result in a directory `DrTutorial/` which contains

- a folder `sequences/` with the SRP input sequences in FASTA format,
- a folder `SHAPE/` with the cotranscriptional SHAPE data,
- a folder `scripts/` with all required `Python` and `R` scripts,
- a folder `drconverters/` with wrappers for various (kinetic) RNA folding methods that convert their output into a common file format, and
- a few additional files with further meta information, such as `README.md` for an overview and description of the repository contents and `pyproject.toml` for installation.

The scripts can be installed from within the `DrTutorial` directory via `pip`:

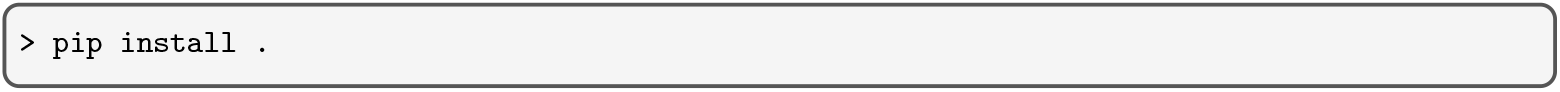

See Note 2 if installation is not desired or impossible.

### 3.4 An overview of computational methods

All of the prediction methods below use the same energy model, the `ViennaRNA` implementation of the nearest neighbor parameters [17, 3]. This enables us to focus on kinetic cotranscriptional effects, rather than subtle differences in energy models for structure prediction.

#### 3.4.1 Minimum free energy (MFE) folding

The MFE can be calculated using `RNAfold` from the `ViennaRNA Package`.

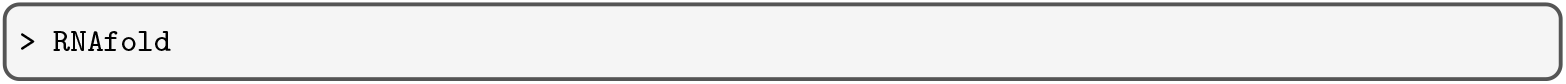

The MFE is of particular interest as it is the thermodynamically most likely structure in the energy landscape. Beware that multiple MFE structures may exist for a specific molecule, see also Note 3. The `RNAfold` program also implements the computation of base-pair probabilities *p*_*ij*_ utilizing McCaskills algorithm [18]. Specifically, the command line option `RNAfold -p` stores the base-pairing probabilities in the form of PostScript dot-plot files (see Note 6).

#### 3.4.2 Sampling the equilibrium distribution

For stochastic sampling from the thermodynamic equilibrium distribution we use `RNAsubopt` from the `ViennaRNA Package`.

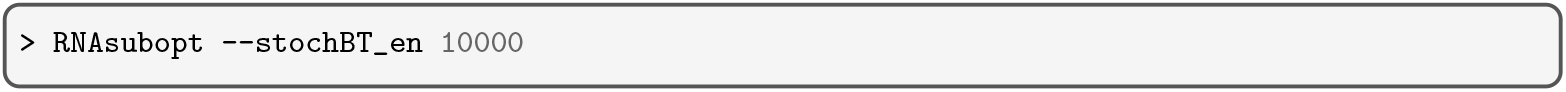

The default mode of `RNAsubopt` is to enumerate all suboptimal structures within a given energy range above the MFE [19]. To obtain a stochastic (Boltzmann) sample from the full equilibrium distribution [20], the option `--stochBT_en 10000` samples 10, 000 structures according to their equilibrium probability and reports their structure, energy and equilibrium probability. See Note 4 for alternative options to obtain stochastic samples.

#### 3.4.3 Sampling the cotranscriptional distribution

For stochastic sampling from dynamically changing cotranscriptional distributions we use the Gillespie-type simulator for nucleic acids `Kinfold`, which assumes that any transition on the secondary structure level must proceed via sequences of “elementary moves”, i.e. either the insertion or deletion of single base-pairs [4].

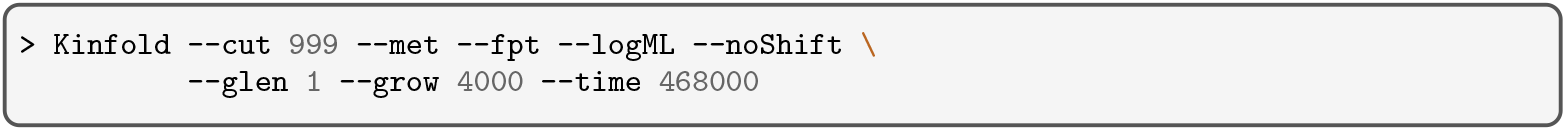

The options (see `Kinfold --help`) are chosen to ensure compatibility with `RNAfold` predictions, and to return a detailed single cotranscriptional folding trajectory using the Metropolis rate model

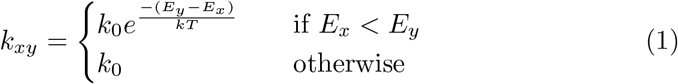

where *k*_0_ is a rate parameter which is scaled by the free energy differences between starting structure *x* and target structure *y*. While we discuss appropriate default values for *k*_0_ below, `Kinfold` only uses free energy differences to calculate rates (i.e. *k*_0_ = 1) and thus expects users to specify the time for nucleotide extensions in internal simulation time units. Here, a nucleotide is attached every 4000 internal time units, and the simulation is set to stop after 117 · 4000 time units, because the molecule is 117 nt long.

For the analysis in this book chapter we provide the script `DrKinfold` (from the `drconverters` repository, see Sec. 3.3) that automatically aggregates data from multiple `Kinfold` trajectories and saves only selected time points.

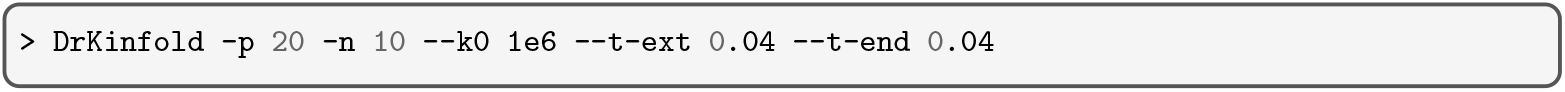

The options `-p 20 -n 10` are chosen to aggregate data from 20 independent `Kinfold` processes, each simulating 10 trajectories. Generally, the options shown above for `Kinfold` are the default options used by `DrKinfold`. However, the options adjusting the rate model use the parameter `--k0`, which translates internal simulation time units into seconds (see Eq. 1), and two parameters `--t-ext` and `--t-end` to set the simulation times during transcription and at the end of transcription in s/nt. For example, the `DrKinfold` call increased the internal simulation time units per nucleotide by a factor 10, as 10^6^ · 0.04 = 40000, corresponding to the `Kinfold` option `--grow 40000`.

#### 3.4.4 Heuristic cotranscriptional folding

Some of the results shown below take many `Kinfold` computation hours. For a faster alternative we use `DrTransformer`, which returns a heuristic, deterministic cotranscriptional structure ensemble [5].

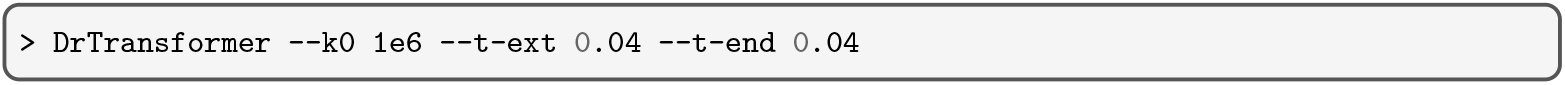

The rate model related command line options of `DrTransformer` are the same as previously introduced for `DrKinfold`, although the rate model is an Arrhenius-type model

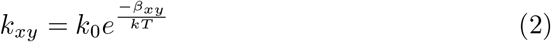

where *β*_*xy*_ is the minimal energy barrier found among elementary paths between starting structure *x* and target structure *y*. (In the `Kinfold` landscape, where all transitions are elementary steps, the two models are equivalent, for more details see [5].) Throughout the chapter we will also use the options `--o-prune` and `--name`, where the former determines how many structures are discarded from the cotranscriptional structure ensemble after the simulation at each nucleotide, and the latter sets the name of `DrTransformer` output files.

### 3.5 A data set of four SRP RNA variants

A drastic example on how difficult it is to determine cotranscriptional folding paths experimentally is the well-studied 4.5S RNA from E. coli which is better known as SRP RNA [21, 22, 23]. Due to the importance of its biological function (guiding and rearranging ribosomes to translocation sites together with SRP proteins), it is known for over 20 years that there are competing metastable cotranscriptional intermediate structures. However, how these structures look like is still under debate [24, 16, 14, 15], and small sequence level variations can have a strong effect on cotranscriptional dynamics [14, 15].

The functional conformation (**FC**) is an extended helical structure interspersed with a few internal loops [25]. It is composed of the hairpin-enclosing helix **H2**, and four stem motifs **S1, S2, S3**, and **S4**, see Fig. 1A. Wong et al. [24] report that a hairpin structure at the 5’ end – **H1a**, see Fig. 1C – is present during two proposed pause sites (U85/U87) but then rearranges into the extended helix.

**Figure 1:**
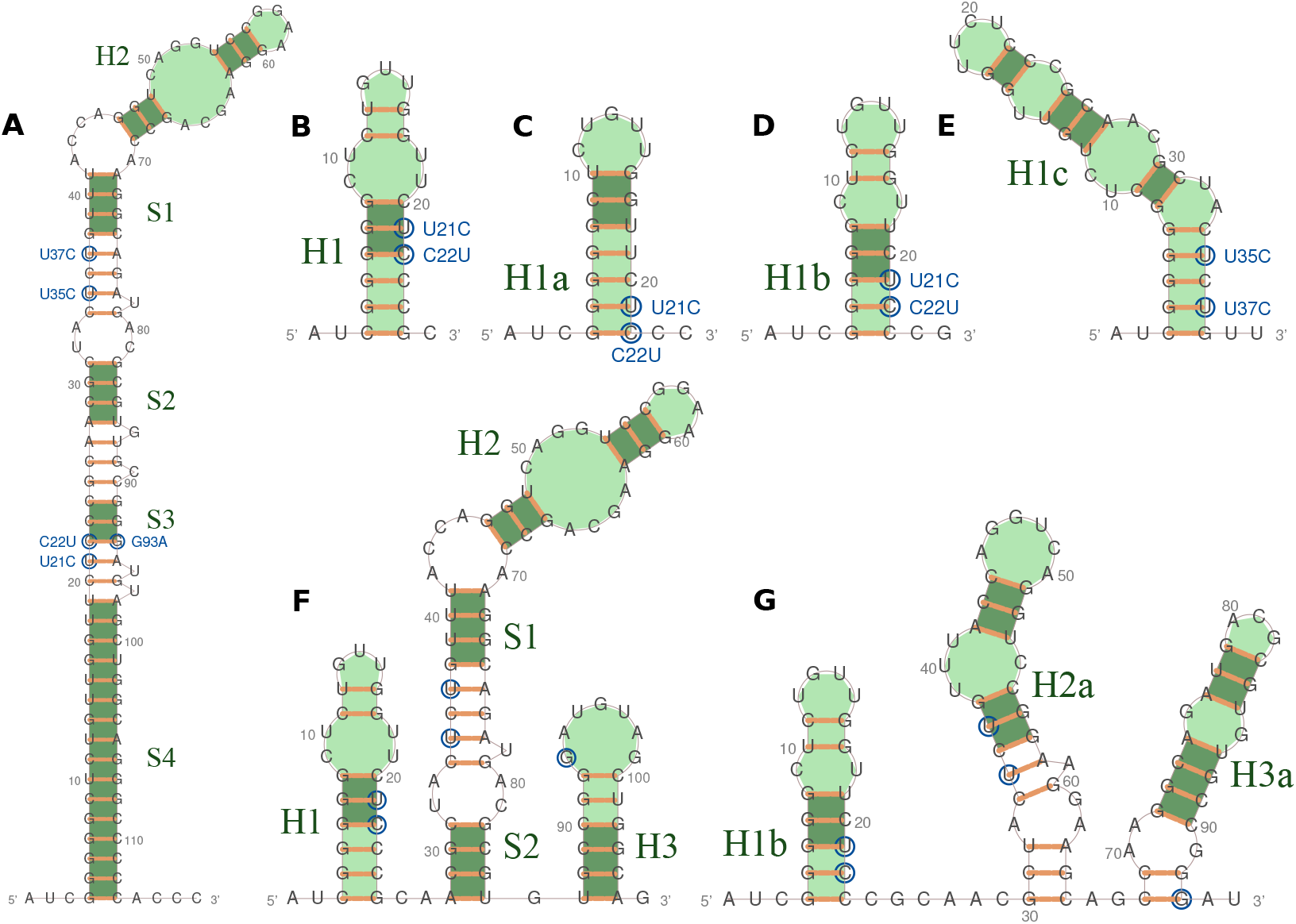
Potential helices formed during SRP transcription. All structures show annotations for the mutations in SRPt (U21C), SRPr (U21C/C22U/G93A), and SRPf (U35C/U37C) indicated by blue circles around the corresponding base. Motif names refer to adjacent regions with green shade; the dark green shade indicates base-pair constraints that define a specific motif in Sec. 3.10. Helix parts not highlighted by dark green color refer to additional compatible base-pairs. (A) The functional conformation (FC) of the SRP RNA. It is a rod-like conformation, consisting of motifs H2, S1, S2, S3 and S4. The mutation U21C destabilizes this conformation, C22U/G93A replace a more stable CG pair with a less stable UA pair. (B) The transient helix H1 modeled after experimental data by Fukuda et al. [14] and Yu et al. [15] is stabilized and destabilized by mutations U21C and C22U, respectively. (C) The transient helix H1a described and experimentally validated by Wong et al. [24] is stabilized and destabilized by mutations U21C and C22U, respectively. (D) The transient helix H1b serving as an alternative conformation to helix H1 and H1a during the rearrangements between H1 and H1a. (E) The transient helix H1c modeled after experimental data by Fukuda et al. [14] is stabilized by the mutations U35C and U37C. (F) An intermediate structure suggested by Fukuda et al. [14] and Yu et al. [15] which is composed of H1, H2, S1, S2 and H3. Note that H2, S1, and S2 are also part of the final functional conformation, while H1 and H3 compete with the formation of S3 and S4. (G) A structure composed of transient helices reported in this tutorial: H1b, H2a, and H3a. H2a is destabilized by U35C and stabilized by U37C. H3a has also been reported as eH3 (early H3) in Yu et al. [15].

Recent cotranscriptional SHAPE-Seq experiments [15] probe the secondary structures at each transcription step to provide additional details on the dynamics: After an initial helix **H1** (see Fig. 1B) has formed, a large part of the molecule forms only small isolated transient helices, until about 109 nt are transcribed. Then, a central helix motif with large interior loops (H2-S1-S2) forms which is also consistent with the final configuration, and a transient helix **H3** forms which encloses the complementary region to the sequence trapped in H1, see Fig. 1F. Interestingly, the authors conclude that the rearrangement between the H1-H2-S1-S2-H3 structure and the final extended H2-S1-S2-S3-S4 (FC) conformation takes place in just a single transcription step from 109 to 110 nt, and suggest a branch migration process via a pseudoknotted intermediate to permit such a fast rearrangement.

Yu et al. [15] also provide additional experiments using point mutations. A sequence with the mutation **U21C** both increases the stability of H1 and destabilizes the functional conformation FC (see Fig. 1). As hypothesized, the U21C mutation prevents the transition into FC by the end of transcription, even though FC is still the thermodynamically favored conformation. A second mutant, **U21C/C22U/G93A**, restores the original stability of H1 while maintaining Watson-Crick complementarity in FC to rescue the cotranscriptional transition into the FC structure.

Recent cotranscriptional optical tweezer experiments [14] identify transcription steps where sudden or continuous folding events take place. The authors independently reported the formation of transient H3 during the transcription of the native SRP sequence, but they also report a transient formation of **H1c** (see Fig. 1E) which competes with H1 before H2 forms. Interestingly, the authors also investigated the U21C mutant, which had the same overall effect as described before, and a second mutant **U35C/U37C** that should stabilize the H1c helix formed early in the transcription process. Cell viability plate assays show that the U21C mutation reduces surviving colony counts, while the U35C/U37C mutations appear indistinguishable.

In this tutorial, we investigate to which extent these experimental findings can be reproduced and predicted *in silico*, but also present new results on the cotranscriptional folding process which – we think – are compatible with the data from experiments. For example, our results neither support the pseudoknot hypothesis from [15], nor do they confirm the H1c structure reported by [14]. Instead, our analysis reveals new insights that involve the intermediate motifs **H1a, H1b, H2a** and **H3a** (see Fig. 1G). We start by defining the following four (partial) SRP RNA sequences taken from Fukuda et al. [14] and Yu et al. [15]:

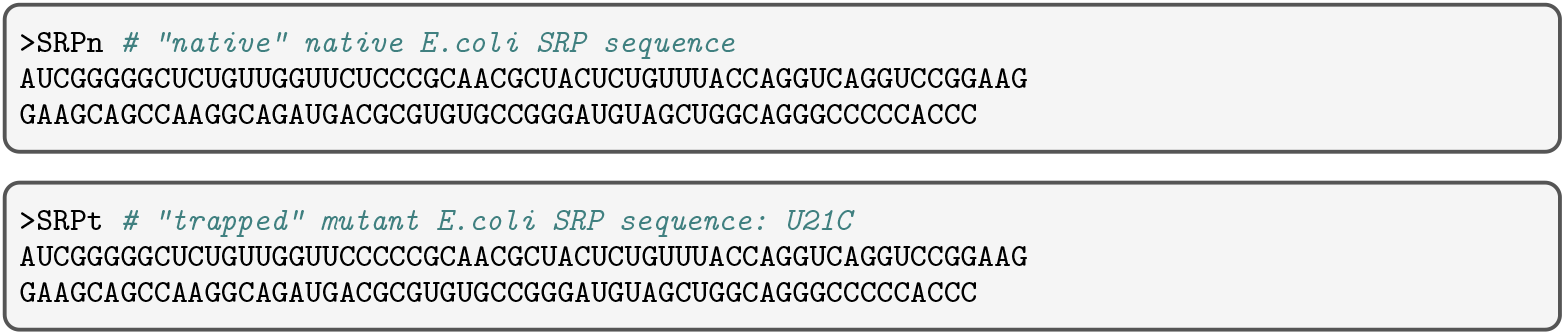
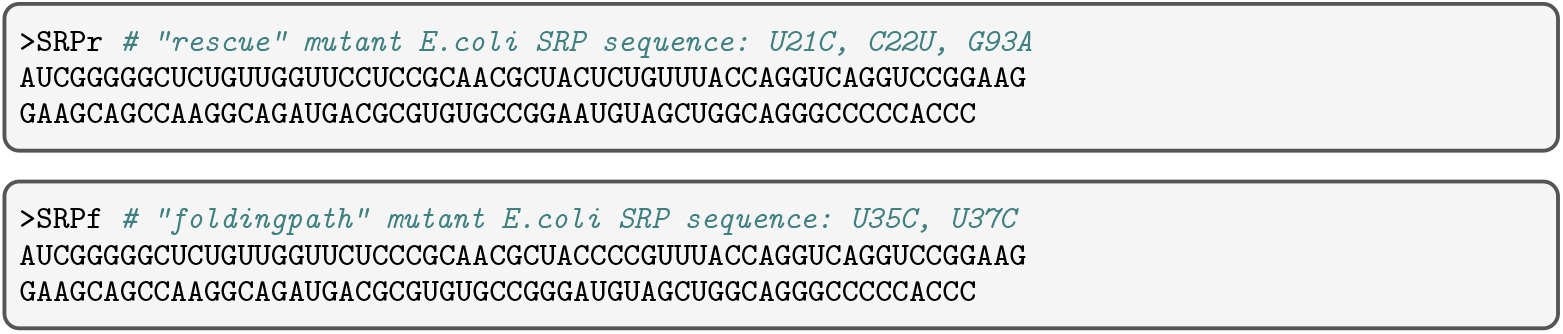

As described in Sec. 3.3 these sequences are also provided in the `sequences/` directory of the `DrTutorial` git repository. They are given the names `SRPn.fa`, `SRPt.fa`, `SRPr.fa`, and `SRPf.fa`, respectively.

### 3.6 Cotranscriptional free energy distributions

As a first step, we aim to identify transcription steps at which the cotranscriptional ensemble is “trapped” and thus differs from the equilibrium distribution. We consider only the free energies of structures for this analysis, as differences in free energy distributions are a simple measure that is always the result of a difference in the underlying structure ensemble. Free energies are also directly related to the probability of observing a structure at equilibrium. Specifically, the probability of any structure *s* is proportional to the Boltzmann factor of its energy *E*(*s*), i.e.

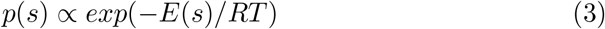

with gas constant *R* and temperature *T* . This implies that finding thermodynamically unlikely structures is difficult using stochastic sampling.

We compare the results of the computational methods `RNAfold`, `RNAsubopt`, `Kinfold`, `DrTransformer` (section 3.4) for each molecule: SRPn, SRPt, SRPr, SRPf (section 3.5). The scripts described below generate a table with 9 columns, which correspond to transcript length, prediction method, the name of the input sequence, the lower and upper 25% quartiles, the median, the mean, the minimum and the maximum values of free energy distributions. The table is stored as a single file in comma separated value (csv) format:

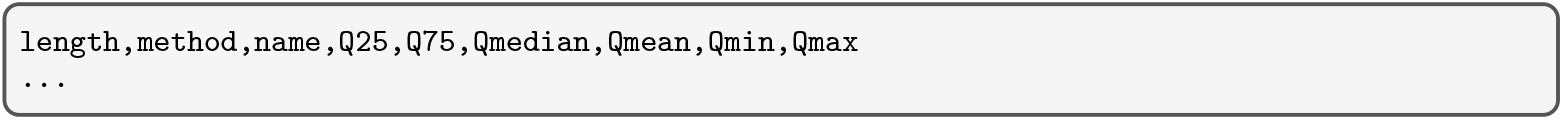

Finally, we plot properties of free energy distributions observed via thermodynamic and cotranscriptional folding methods (see Fig. 2). A similar figure – also for the SRP RNA – can be found in Yu et al. [15].

**Figure 2:**
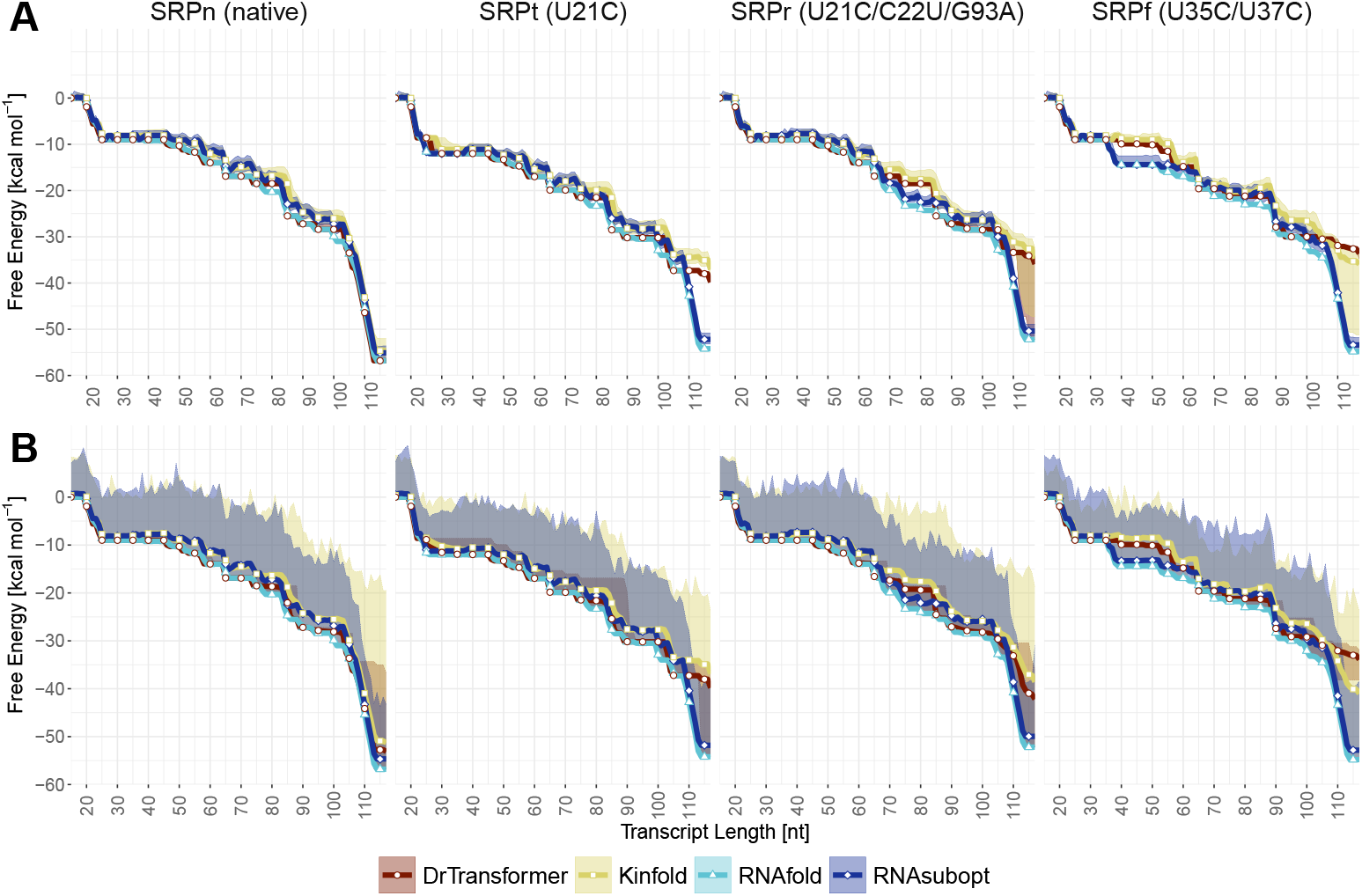
Free energy distributions of thermodynamic and kinetic folding predictions. The four columns compare SRP-WT, SRP-U21C, SRP-U21C-C22U-G93A, and SRP-U35C-U37C. All plots show the MFE (`RNAfold`), 10, 000 stochastic samples from the equilibrium distribution (`RNAsubopt`), 10, 000 stochastic cotranscriptional trajectories (`Kinfold`) and the energy distribution from a heuristic method (`DrTransformer`). **(A)** Thick solid lines denote the median free energy of the prediction, while semi-opaque bands indicate the first and third quartile, respectively. **(B)** Thick solid lines denote the mean free energy of the prediction, while semi-opaque bands indicate the minimum and maximum values, respectively.

#### 3.6.1 Cotranscriptional distributions

To calculate the free energy distributions during cotranscriptional folding, the program `DrKinfold` is called for each SRP variant:

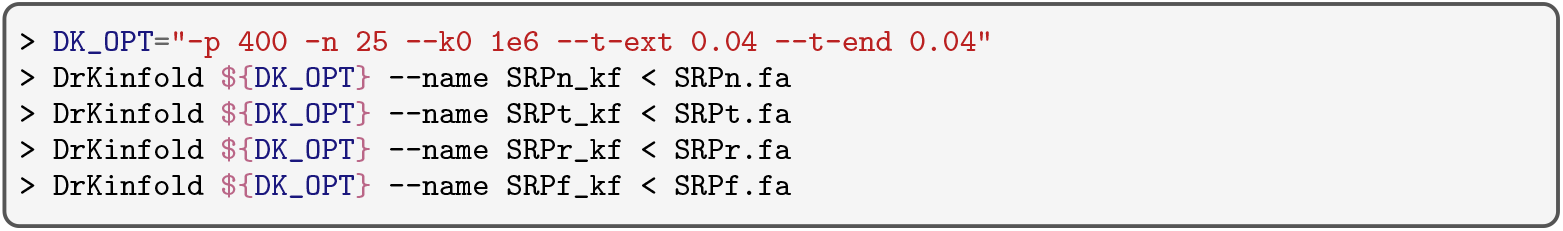

Here, distributions are calculated from 400· 25 = 10, 000 trajectories to get a broad distribution of min/max values. (We recommend using 100 simulations for testing, as this is sufficient to reproduce most of the important results.) The options `--k0 1e6`, `--t-ext 0.04`, and `--t-end 0.04` adjust the kinetic model.

Also the program `DrTransformer` is called for each SRP variant:

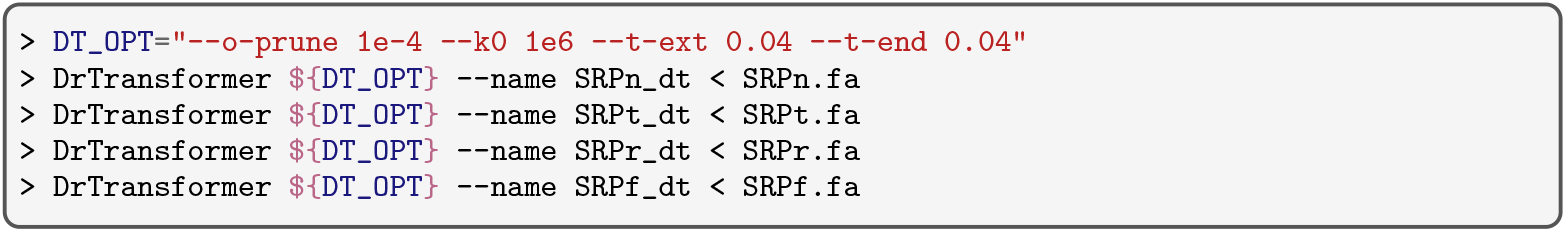

Here, `--o-prune 1e-4` is used as a cutoff to discard low occupied secondary structures after each simulation and the remaining options adjust the kinetic model.

Both `DrKinfold` and `DrTransformer` produce `{--name}.drf` files, which contain a table with the output of simulations. The file suffix `.drf` denotes that these files are suitable input for the visualization software `DrForna`, which we will use in Sec. 3.8. We provide a parser script `drf_parser.py` to extract the distributions of free energies at the end of the simulation for each nucleotide, i.e. before the next nucleotide is appended during transcription. The following commands perform these computations

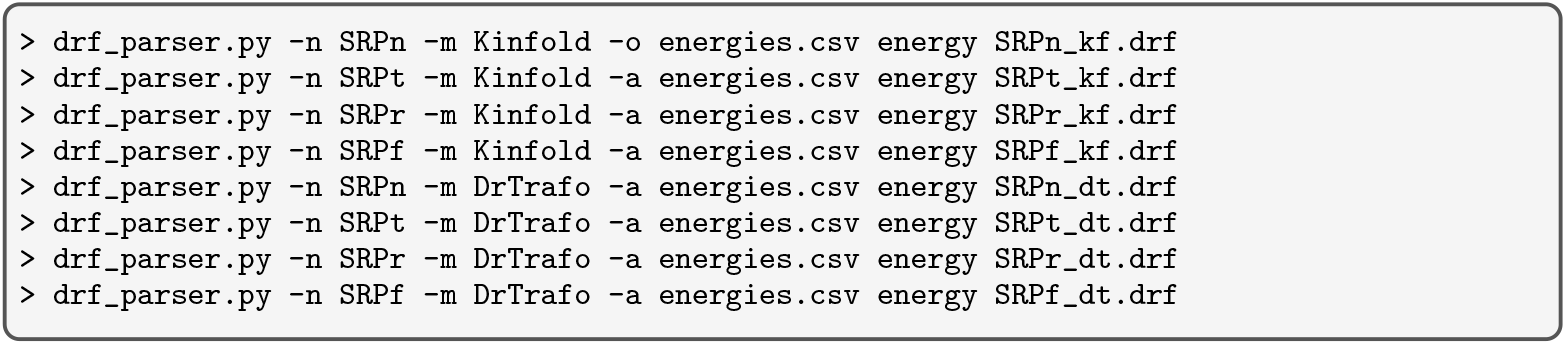

where for each of the four input drf files the script is run in *energy distribution mode* (keyword: `energy`) and the options (`-n -m`) set the name and the method fields for the csv file. Note the option `-o energies.csv` to write a new output file which also includes the header information, and the option `-a energies.csv` to append the output to an existing file without the header.

#### 3.6.2 Thermodynamic distributions

To compare the cotranscriptional folding effects with a thermodynamic approach, the equilibrium ensembles have to be calculated for each intermediate length of the growing, nascent RNAs. The procedure is simple:

1. Read the full length input sequences
2. Loop over subsequences for all transcript lengths
3. Predict the MFE for the subsequences
4. Perform stochastic sampling for the subsequences.

Instead of calling the command line executables `RNAfold` and `RNAsubopt`, we use the `ViennaRNA Package` Python interface (via the 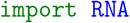 Python statement) to write a script that implements the full procedure stated above:

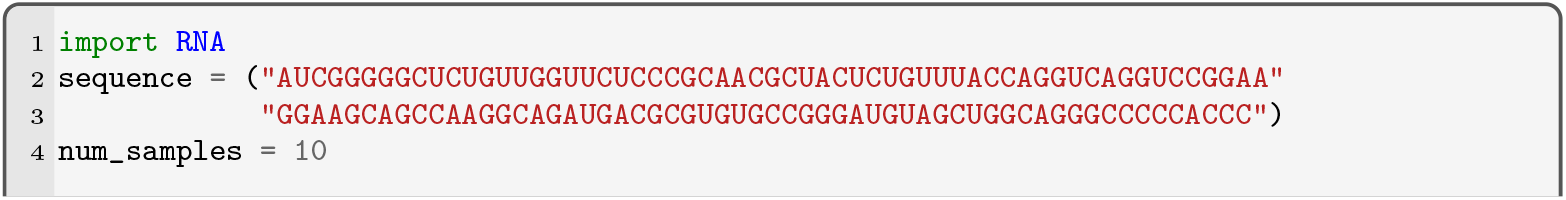
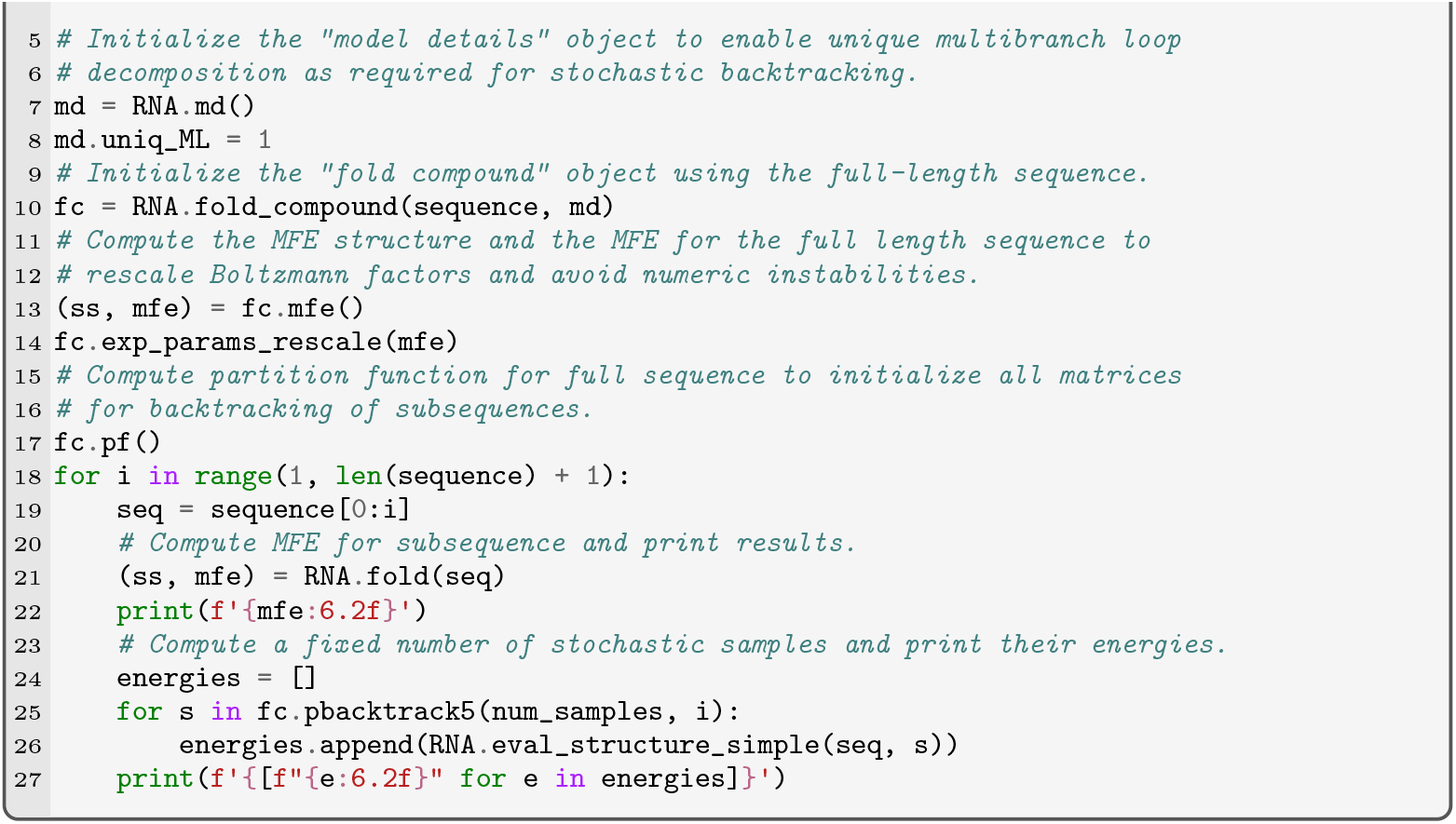

Note, that the above code is a simplified version of the code implemented in the script `thermo_predict.py` that is provided in our git repository. The latter provides various additional options and prints results in the correct output format. To compute the thermodynamic energy profiles with our `RNAfold`/`RNAsubopt` wrapper script use the following commands:

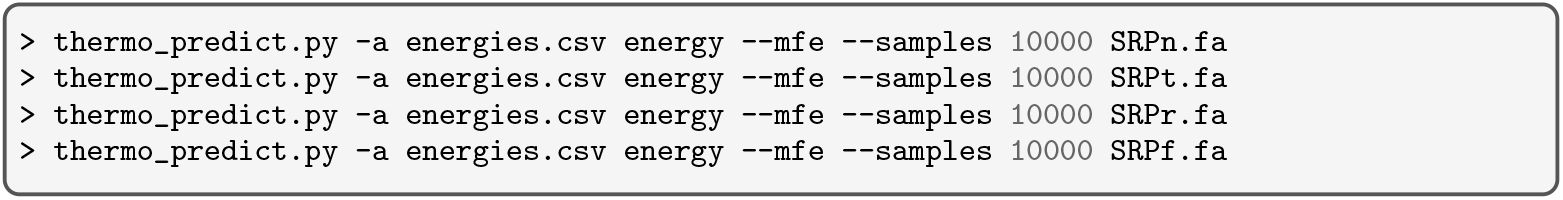

For each of the four SRP input sequences the script appends (`-a`) data to the file `energies.csv`. The keyword `energy` runs the script in *energy distribution mode* to compute both the MFE (`--mfe`) and the observed distributions from 10, 000 stochastic samples (`--samples 10000`).

#### 3.6.3 Interpreting the results

Having all energy distributions stored in the file `energies.csv` allows us to investigate the differences between thermodynamic and kinetic folding behavior. We plot the data using an `R` script provided in our git repository:

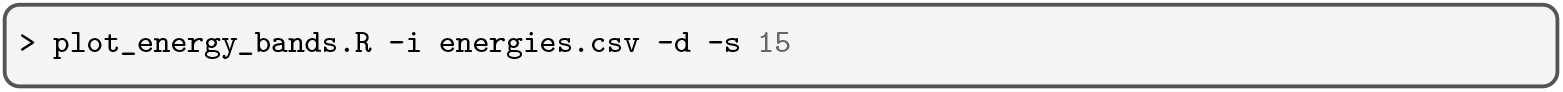

Here, we use the `-i energies.csv` option to specify the input file, and the options `-s 15` and `-d` to start the plot with transcripts of length 15. Fig. 2 shows two plots for each SRP variant: the top row depicting the median energies together with the range between the 25% and 75% quartiles *Q*_[25,75]_ (Fig. 2A), and the bottom row showing the mean energies and the corresponding minimum and maximum values (Fig. 2B).

Generally, we compare `RNAfold` with `DrTransformer` predictions, and `RNAsubopt` with `Kinfold` samples. `DrTransformer` only considers local minima during a simulation, and the MFE returned by `RNAfold` is always the most dominant minimum in the equilibrated ensemble. In contrast, `RNAsubopt` and `Kinfold` both sample structures from the full secondary structure ensemble, which means the mean and median of those samples must be higher than for the other methods. For the sequence “native SRPn” the results of cotranscriptional simulations are mostly comparable with thermodynamic approaches. The medians and the *Q*_[25,75]_ range (Fig. 2A) are similar for `RNAfold` and `DrTransformer` results, as well as for `RNAsubopt` and `Kinfold` results. Notably, around 80 and 100 nt the `DrTransformer` simulations suggest that the MFE structure does not dominate the ensemble. This can also be seen when comparing `RNAsubopt` samples with `Kinfold` samples, especially when looking at the mean energies and the min/max range (Fig. 2B). Also, we observe that there is a point around 110 nt length, where the majority of structures is still in an energy range close to the MFE, but cotranscriptional simulations also start observing structures in a much higher energy range, compared to the thermodynamic approaches.

The mutated sequence “trapped SRPt” (mutation U21C) has an altered MFE profile during transcription: the mutation lowers the MFE after about 25 nt have been transcribed, and this effect persists until roughly 105 nt have been transcribed, then the effect is reversed, and the MFE is higher than that of the native SRP molecule. Interestingly, trapped SRPt shows the same differences between thermodynamic and cotranscriptional approaches around 80 and 100 nt that we already observed for the native SRPn molecule, indicating that the mutation has no effect on those cotranscriptional folding events. After 105 nt, the cotranscriptional simulations clearly diverge from thermodynamic predictions, as the ensemble is trapped about 15 kcal · mol^−1^ above the MFE, and this effect persists until the full molecule is transcribed. However, even though most samples are trapped, both `Kinfold` and `DrTransformer` observe structures in the range of the MFE.

The mutated sequence “rescue SRPr” (mutations U21C/C22U/G93A) restores the native SRPn MFE profile until about 65 nt have been transcribed. Then, cotranscriptional simulations stay in an energy regime about 5 kcal · mol^−1^ above the thermodynamic predictions roughly until nucleotide 85 is transcribed. Eventually, we observe a similar effect as with SRPt: after 105 nt the majority of cotranscriptional folds is trapped about 15 kcal · mol^−1^ above the MFE. In contrast to SRPt, the ensemble distribution at the end of transcription is much more diverse as the MFE is now within *Q*_[25,75]_ of both cotranscriptional methods. This indicates that some molecules fold into the MFE structure.

The mutated sequence “foldingpath SRPf” (mutations U35C/U37C) shows strong cotranscriptional folding effects already after 35 nt have been transcribed. The sequence mutations have lowered the MFE and equilibrium distribution between 35 and 65 nt, but the cotranscriptional ensembles do not follow this trend. Comparisons to the other molecules are difficult, because neither the thermodynamic nor the cotranscriptional ensembles follow the same patterns as before. Perhaps surprisingly, between 80 and 90 nt we observe an increase of cotranscriptional ensemble free energies. This may point to a degenerate landscape with many competing structures of similar free energy. Between 90 and 100 nt `DrTransformer` suggests that most of the ensemble is equilibrated, while `Kinfold` ensembles are trapped at a higher energy range. This effect is inverted at 105 nt, when both the `Kinfold` and `DrTransformer` ensemble are trapped in high energy conformations, but `Kinfold` has a much broader distribution that also contains the MFE configuration.

### 3.7 Cotranscriptional accessibility profiles

The previous analysis successfully identified cotranscriptional traps in the folding process, but we have no indication on what structures are actually involved. This next analysis is a first step toward visualizing the different structures that molecules adopt during transcription. Specifically, we plot the probability of being (un)paired for each nucleotide. This measure can be plotted in one dimension (as opposed to base-pair probabilities which require a two dimensional representation, see Note 5) and it is also useful to relate structure prediction with RNA structure probing methods such as cotranscriptional SHAPE [26], which we will address later in Sec. 3.9. Visually observing how (un)pairedness changes during transcription can reveal structural rearrangements and thus implicitly shows the transitions between dominant secondary structure motifs. The result of this analysis is shown in Fig. 3.

**Figure 3:**
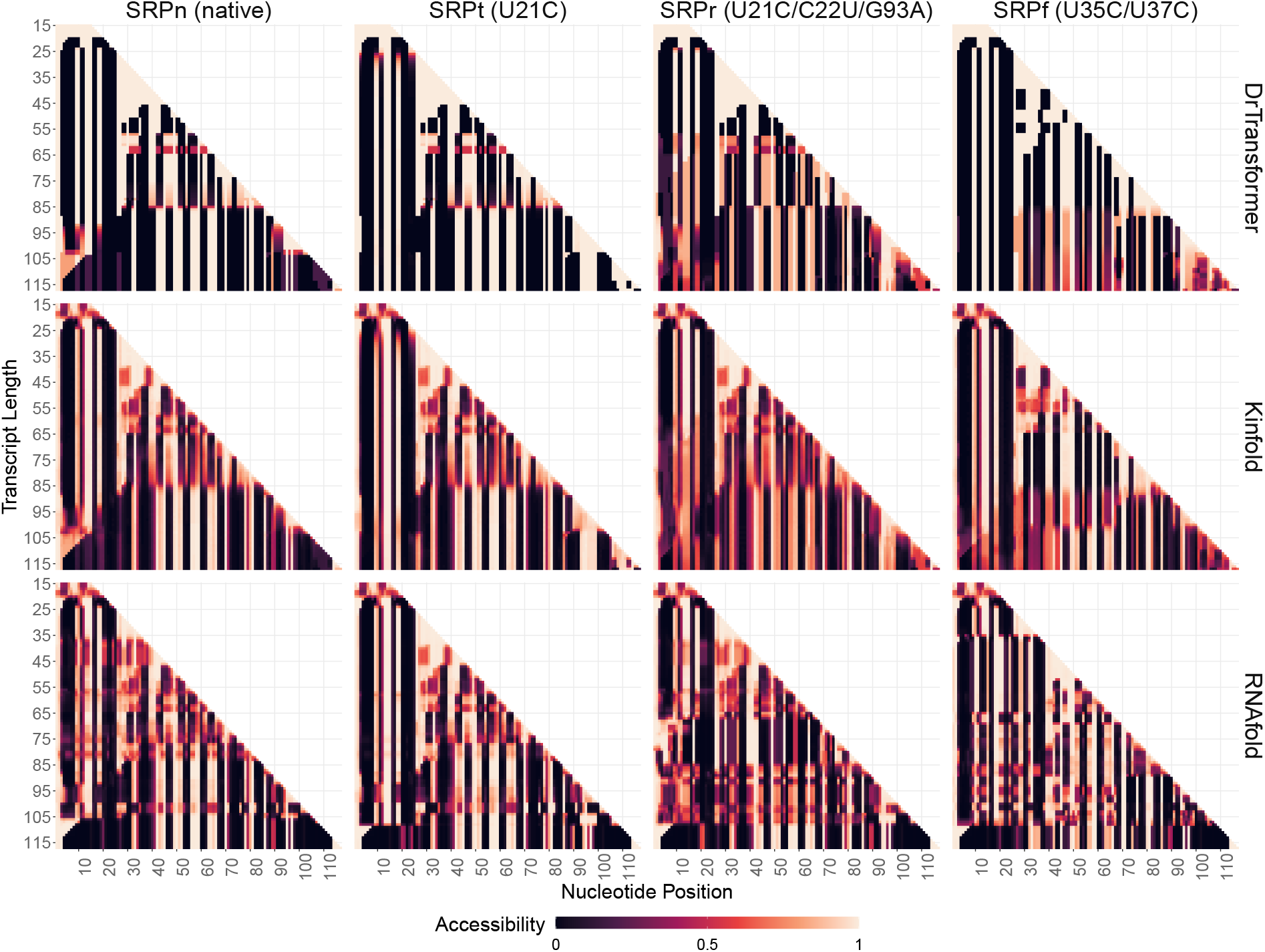
Cotranscriptional accessibility profiles. The change of (un)pairedness is shown as a function of transcript length for four different SRP sequences and three different computational models.

Instead of calculating the probability that nucleotide *i* forms a base-pair with any *j*, we calculate the opposite: that *i* does not form any base pair. In literature the latter is often called **accessibility**, indicating that the nucleotides are available for interactions with the environment. Formally, we can compute the accessibility *q*_*i*_ of nucleotide *i* as

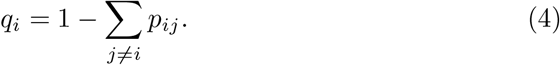

where *p*_*ij*_ is the equilibrium probability of a base-pair (*i, j*) between two compatible nucleotides *i* and *j*. The equilibrium probability *p*_*ij*_ can be computed from the set of structures *S*_*ij*_ = *{s* | (*i, j*) ∈ *s}* that contain (*i, j*) and the partition function *Z* = ∑_*s*_ *exp*(−*E*(*s*)*/RT*), as

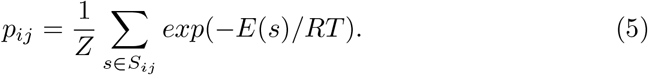

For kinetic folding simulations, we calculate the relative frequencies *f*_*ij*_ of observing base-pair (*i, j*) among all structures present at a specific time point

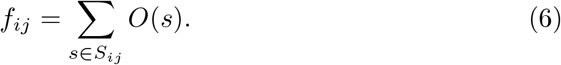

where *O*(*s*) is the “occupancy” of a structure – the probability of observing a structure in a kinetic simulation. The occupancy can be derived from a stochastic simulation as 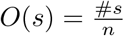 where #*s* is the number of times structure *s* is observed in *n* samples at a specific time point, or it can be calculated numerically in the deterministic models.

#### 3.7.1 Cotranscriptional accessibility profiles

We use the *.drf output files from `Kinfold` and `DrTransformer` as produced in Sec. 3.6.1 to extract the accessibility of each nucleotide during the cotranscriptional folding process. The script `drf_parser.py` in *accessibiilty calculation mode* (keyword: `accessibility`) derives accessibilities from structure ensembles at the end of the simulation for each nucleotide, i.e. before the next nucleotide is transcribed.

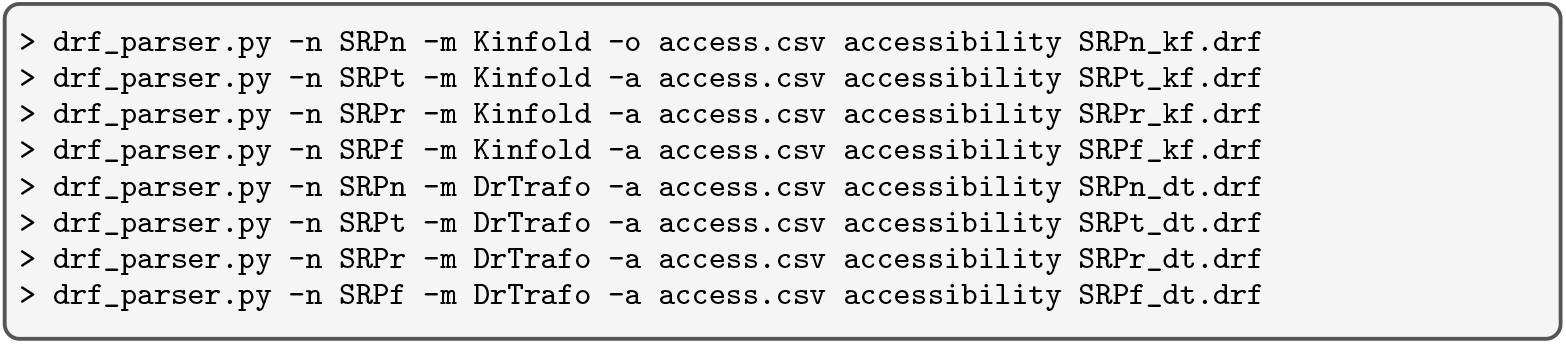

Note the similarity to respective command line calls discussed in Sec. 3.6.1. Here, the relevant data is stored in the comma separated value file `access.csv`.

#### 3.7.2 Thermodynamic accessibility profiles

We compare the cotranscriptional with thermodynamic accessibility profiles, where the latter have to be calculated for each intermediate length of the growing, nascent RNAs. As before, the `thermo_predict.py` script uses the `ViennaRNA Package Python` API to perform all necessary computations. For each transcript length *l*, we compute the base-pairing probabilities 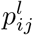 to derive 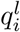 according to Eq. 4. The code block required for this task is as follows:

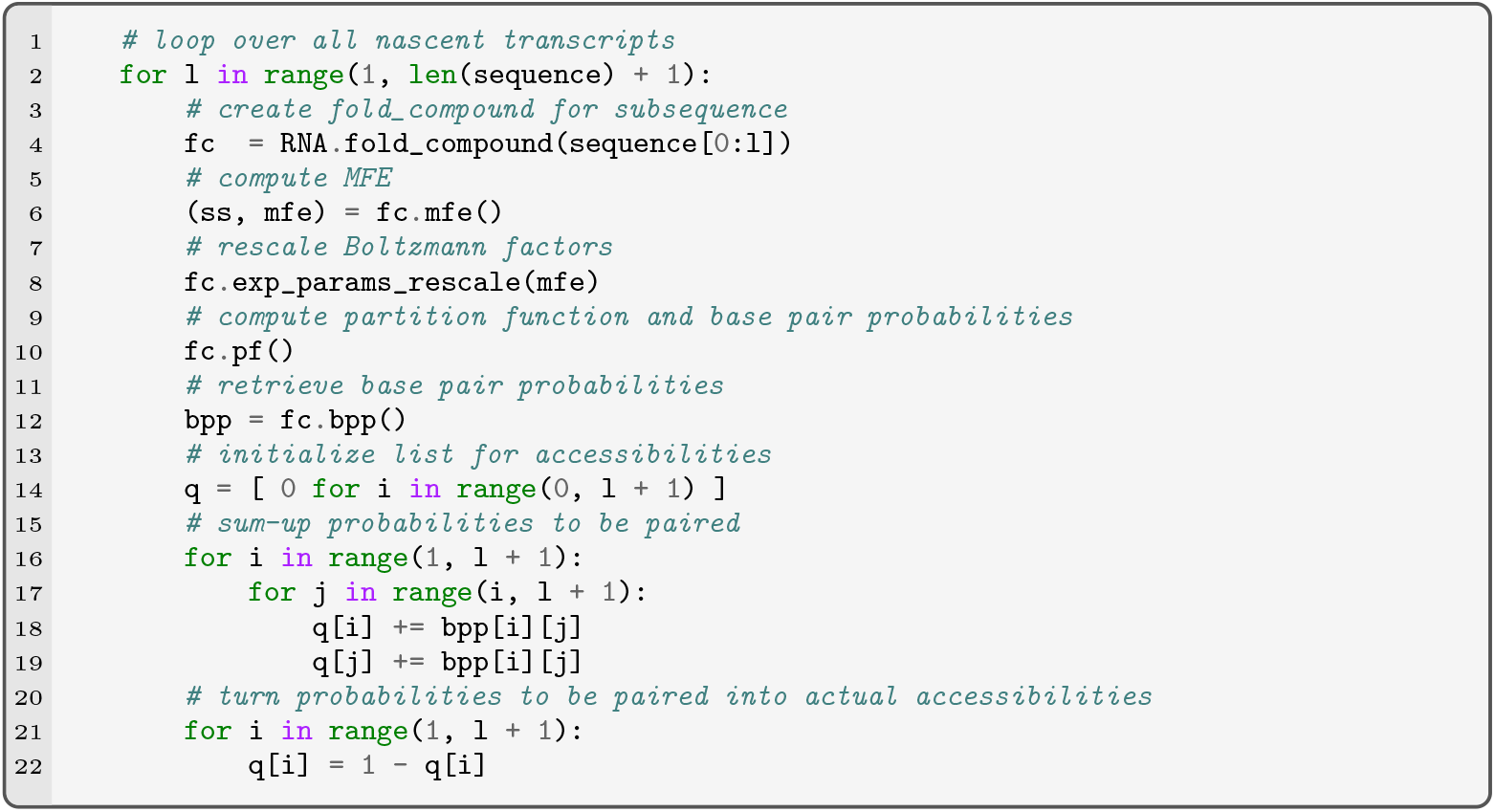

The script `thermo_predict.py` in *accessibiilty calculation mode* (keyword: `accessibility`) uses this code to calculate thermodynamic accessibilty profiles:

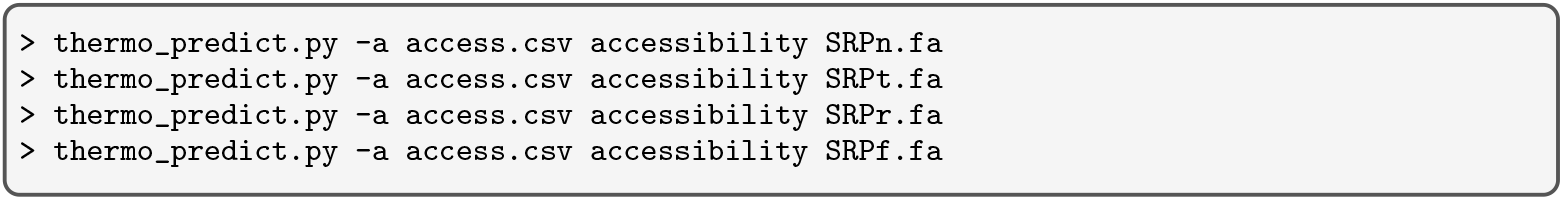

As before, the above commands append the resulting accessibilities 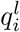 for all four SRP RNA sequences to the already existing `access.csv` file.

#### 3.7.3 Plotting and interpreting the results

We use the data from the generated file `access.csv` together with the script `plot_accessibility.R` from the git repository to observe the changes in accessibility as the RNA molecule is transcribed (see Fig. 3).

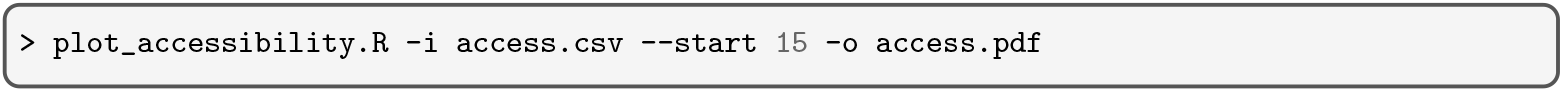

`DrTransformer` predicts less variance in accessibility than `Kinfold` and `RNAfold` across all heatmaps of Fig. 3. This is expected, as the (un)pairedness for `RNAfold` and `Kinfold` is computed from the entire structure ensemble and a large amount of folding trajectories, respectively. In contrast, `DrTransformer` considers a relatively small number of local minimum structures during a simulation. While the information of what is paired with what is not explicitly shown, symmetric patterns of pairedness (dark areas) typically correspond to regions pared with each other. Note that it can be helpful to compare the results here with the results from Fig. 2 as the information about energy distributions and structure ensembles together gives a much more complete impression of the underlying folding events than each method by itself.

For the native SRPn RNA sequence, `DrTransformer`, `Kinfold` and `RNAfold` provide similar results. Following `DrTransformer` results, the first stable H1 helix forms immediately after the first 25 nt have been transcribed. A second, temporary helical structure forms between 45 and 55 nt, at 58 nt that structure transitions into a new intermediate which builds up until approximately 84 nt have been transcribed. Finally, at 85 nt the H2 hairpin of the final functional conformation forms and extends gradually to eventually displace H1. Note, that there are competing structures, e.g. at 105 nt H1 is not dominant, but the shades of red indicate that there is still a fraction of H1 paired in the ensemble. From the results in Fig. 2 we expect differences between the cotranscriptional and thermodynamic predictions at 80 and 100 nt. Indeed those are regions where `RNAfold` shows different patterns of pairedness. At 80 nt, the H2 helix is already dominant in the thermodynamic ensemble, while cotranscriptional simulations are still trapped in the H2a intermediate structure. At transcript lengths of about 100 nt, `RNAfold` suggests a sharp temporary transition to an ensemble with an alternative accessibility profile, but this temporary ensemble is not relevant during cotranscriptional simulations. Interestingly, `RNAfold` also hints toward an alternative structure between 36 and 45 nt which competes with H1, but does not play a role in `Kinfold` and `DrTransformer` simulations.

For the trapped SRPt RNA sequence, `RNAfold` predictions are similar to those of the native SRPn, but one can see that the mutation stabilizes the H1 helix, as less alternative structures are visible. The `DrTransformer` and `Kinfold` simulations, however, show that the RNA is cotranscriptionally trapped in structures that contain the H1 helix. While this does not effect the formation of the H2 seed (which happens at the same time as in SRPn), H1 never gets removed and instead, the final structure contains H1, H2 and a new H3 conformation, which forms after 105 nt have been transcribed. (A faint indication of H3 forming can also be seen in the cotranscriptional simulations of SRPn.)

The accessibilities for the rescue mutant SRPr show that for this RNA many more structural alternatives exist during transcription. `DrTransformer` simulations are similar to SRPn and SRPt sequences up to about 90 nt, except for the faint red background indicating a larger variety of structures in the ensemble. After 90 nt, `DrTransformer` show a mix of the SRPn and SRPt folding paths, suggesting that H1 is being replaced by some structures that (presumably) form the final functional conformation (H2-S1-S2-S3-S4). Like for SRPn and SRPt, the `Kinfold` data is a noisy version of the `DrTransformer` results. The `RNAfold` data, however, looks different from cotranscriptional simulations. Between lengths 65 and 85 nt equilibrium conformations do not contain the H1 helix, which seems to explain the large difference of free energy distributions observed in Fig. 2. After 85 nt we see more similarities with cotranscriptional data, and – in combination with results from Fig. 2 – conclude that the cotranscriptional ensemble contains dominand structures from the equilibrium ensemble.

The last column in Fig. 3 shows the accessibilities of the foldingpath mutant SRPf. We see differences in all three prediction methods `DrTransformer`, `Kinfold`, `RNAfold`. As originally proposed by Fukuda et al. [14] the mutations are meant to stabilize H1c, a helix competing with H1 even before H2 is forming (H2 forms typically around 65 nt). This is clearly visible in the `RNAfold` predictions at about 38 nt length (cf. SRPn), which suggest that this new helix is dominant until about 110 nt. The mutations also favor the formation of the initial H2 hairpin earlier than in other molecules. However, these results are drastically different from cotranscriptional simulations. `Kinfold` and `DrTransformer` agree that H1 is dominant throughout transcription, and that the mutations stabilize a different intermediate structure between 38 and 45 nt length which is compatible with H1. After about 58 nt, the second intermediate structure (H2a) is stabilized. After 85 nt, `DrTransformer` and `Kinfold` predict opposing folding events. According to `DrTransformer` H2a is dominant until the end of transcription, although we see a faint background that indicates H2 seed formation. In contrast, `Kinfold` suggests a transition into the functional structure (H2-S1-S2-S3-S4) at around 105 nt. Thus, `Kinfold` results suggest that a part of the ensemble reaches the functional structure at transcription end, while `DrTransformer` predicts the ensemble to be trapped in a non-functional form.

### 3.8 Visual inspection of the structures

The analyses in Fig. 2 and Fig. 3 provide a high level overview on how ensembles change during transcription. We can use `DrForna` to see whether those changes correspond to structures that have been previously described in literature (see Fig. 1), or to novel conformations. `DrForna` is a Javascript application that displays the secondary structures of cotranscriptionally changing ensembles. While we encourage users to visit the `DrForna` project ^1^ and load the respective *.drf files generated in this tutorial, we show a few selected transcription steps for the native SRPn and `DrTransformer` results in Fig. 4.

**Figure 4:**
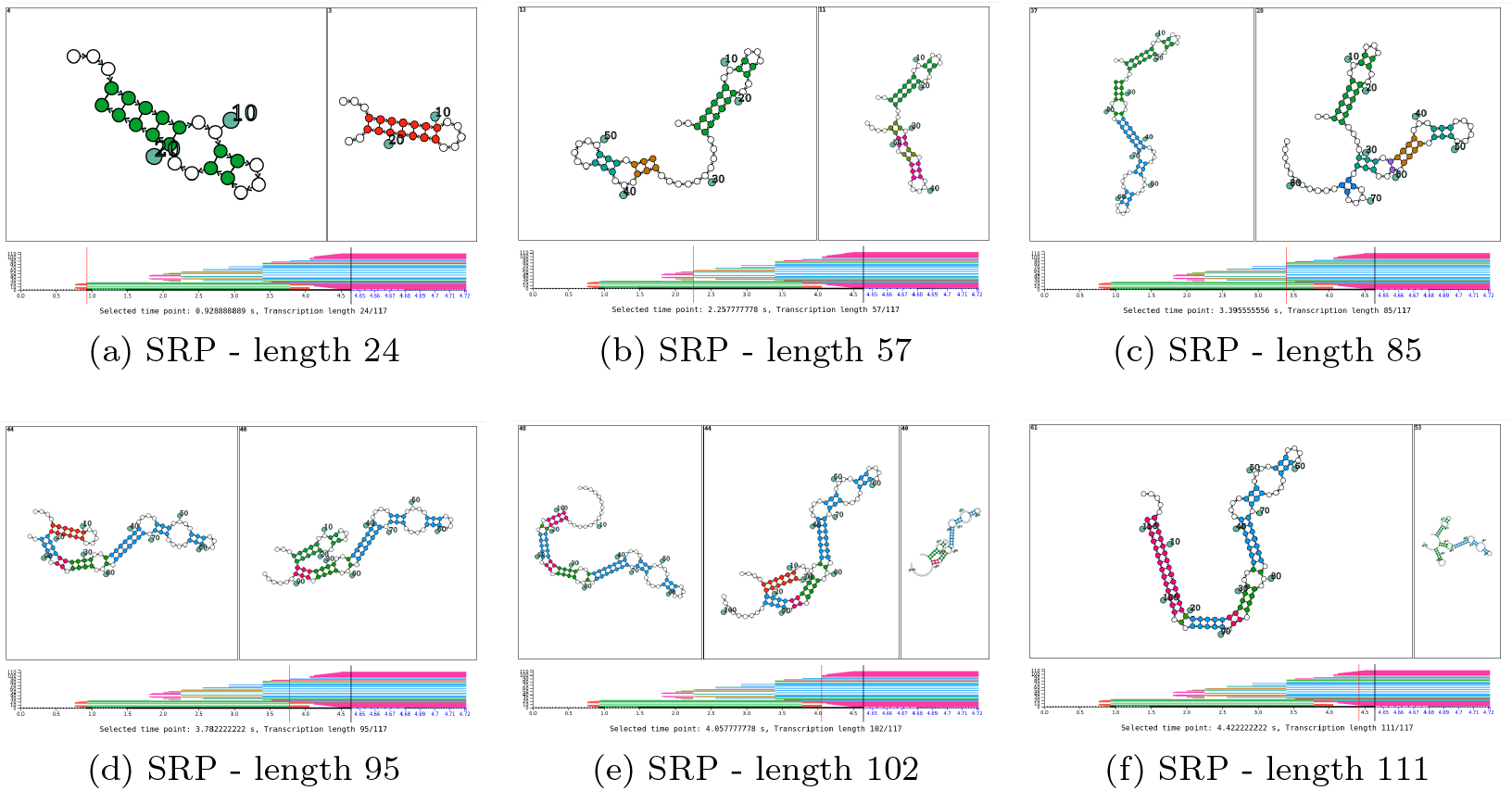
Visual inspection of selected structure transitions.

Fig. 4(a) shows SRPn at 24 nt: H1a transitions into H1. Fig. 4(b) shows SRPn at 57 nt: the first (unnamed) intermediate structure transitions into H2a. Fig. 4(c) shows SRPn at 85 nt: H2a transitions into H2-S1-S2. Fig. 4(d) shows SRPn at 95 nt: While the S3 stem is forming, there is another competition between H1 and H1a (cf. Fig. 4(a)). Fig. 4(e) shows SRPn at 102 nt: The competition from the previous panel continues, but now a third structure is dominating the ensemble where S4 is (partially) formed, and neither H1 nor H1a are present. Fig. 4(f) shows SRPn at 111 nt: The functional conformation H2-S1-S2-S3-S4 is in competition with the H1-H2-S1-S2-H3 structure that was observed by Yu et al. [15].

A key moment revealed by visual analysis is the competition between H1 and H1a, which is not only relevant during the early formation of H1, but also during the formation of the S3 stem of the final functional structure. To learn about the transition landscape between H1 and H1a in more detail, we show a screenshot of `Kinfold` results visualized with `DrForna` in Fig. 5. It shows a short lived intermediate conformation H1b, which may be relevant during the the transition from H1 to H1a. This rearrangement provides an alternative hypothesis on how the final H2-S1-S2-S3-S4 conformation can form in such a short time, without postulating a much more complex reconfiguration via a pseudoknotted intermediate (cf. Yu et al. [15]).

**Figure 5:**
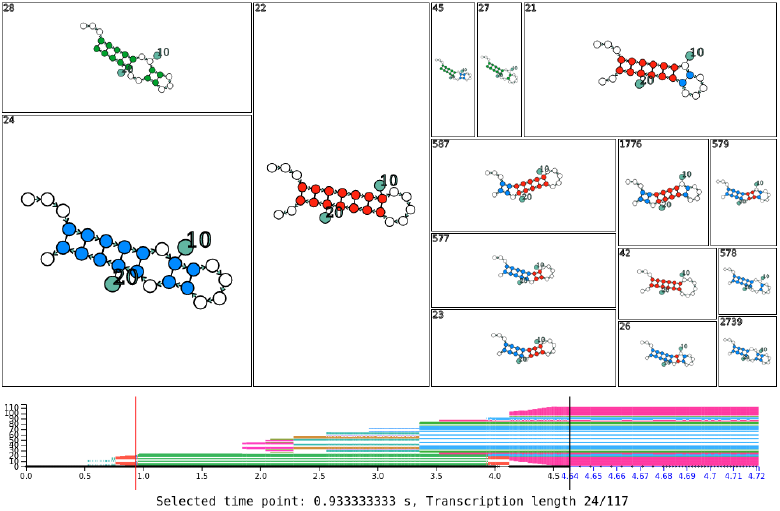
A `DrForna` snapshot to learn about the transition landscape from H1a (red) to H1 (green) in detail. The analysis reveals a new transient hairpin H1b (blue) which can be generated from both H1a and H1 via a moving bulge loop.

### 3.9 A comparison with experimental data

As the analysis provided in this contribution is inspired from experimental work on the SRP RNA [24, 16, 14, 15], we provide a high-level comparison between experimental and simulated data on SRP RNA folding. To that end, we use SHAPE-Seq data provided by Watters et al. [16] and Yu et al. [15] and then modify the cotranscriptional model parameters such that they are closer to the experimental SHAPE-Seq setup. The results are compared visually in Fig. 6. A more detailed analysis of experimental data in combination with cotranscriptional predictions is beyond the scope of this contribution, but we hope that this approach may inspire further work in this direction.

**Figure 6:**
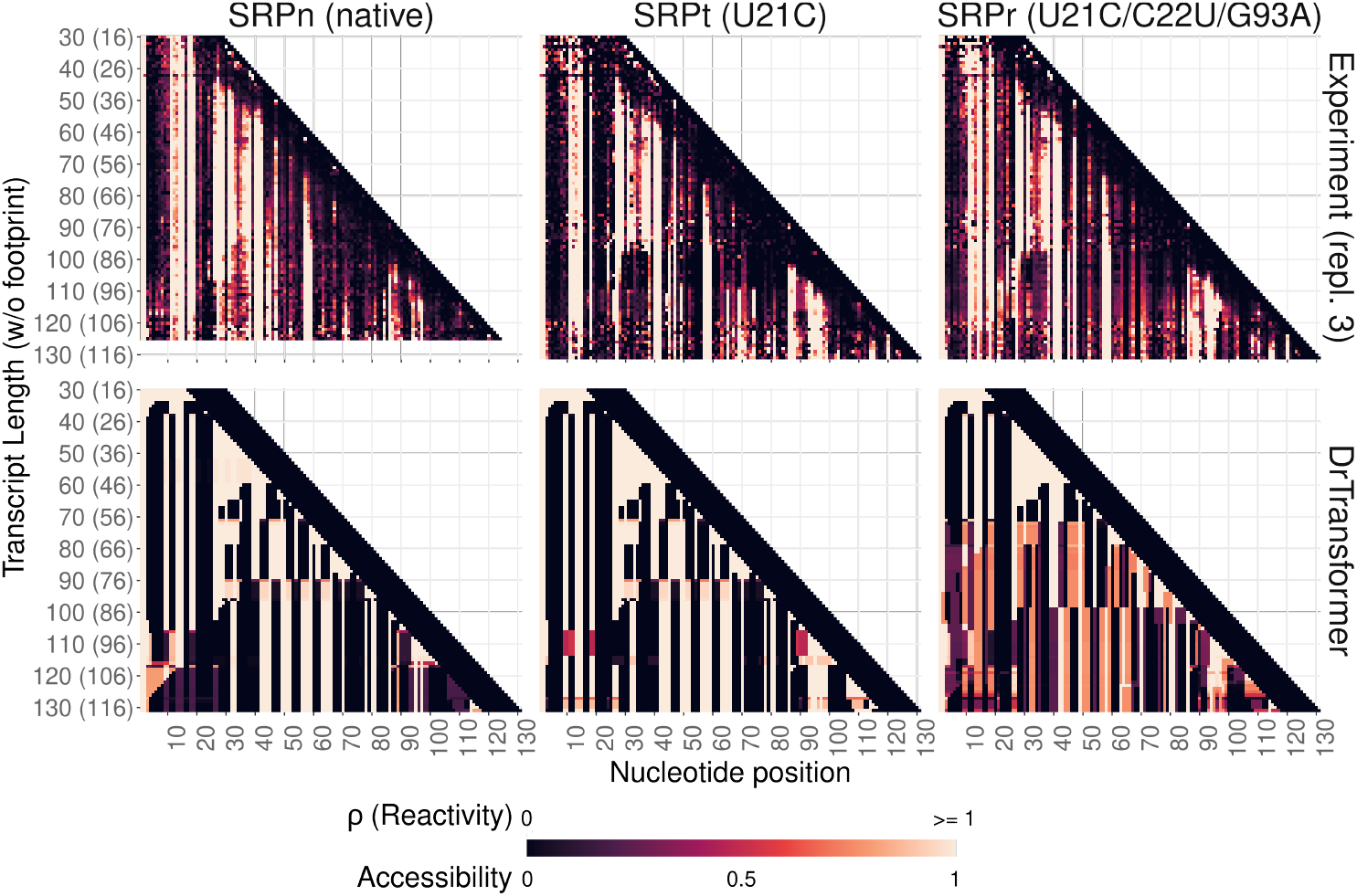
Comparison against cotranscriptional SHAPE experiments. Shown in the first row of heatmaps are experimentally derived SHAPE reactivities (replicate 3) for the native SRP sequence SRPn [16], and the two mutants SRPt and SRPr [15]. The second row depicts accessibilities 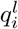 as computed by cotranscriptional folding simulations with `DrTransformer`. Since the experimental protocol allows the nascent RNA sequences to refold for 30 s before they are probed the folding simulations have been adopted accordingly. Although there is no linear relationship between SHAPE reactivities *ρ* and accessibilities 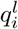, both the experiment and predictions are in generally good agreement with respect to the major helices that are formed. The rightmost diagonal lines with low reactivities (resp. accessibilities) denote the RNA polymerase footprint.

#### 3.9.1 Adjusting the predictions to the experiment

SHAPE measures the accessibility of nucleotides by incorporating a reactive agent into flexible regions of the backbone, e.g. paired bases are much less flexible and thus yield low reactivities. It is well known that reactivity profiles from high-throughput experimental structure probing methods in general do not correlate directly with the probability of being unpaired in the secondary structure model [27, 28, 29, 30]. In part, this observation can be attributed to the fact that SHAPE probing also reveals tertiary interactions instead of just secondary structure, but also other known or yet unknown biases influence the reactivities. For example, potential sequence dependence of the reagent, unusually flexible bases at the end of helices or inflexible nucleotides forming non-canonical base-pairs within seemingly unpaired regions, as well as biases in the relative positioning toward the 3’ vs 5’ end [31, 32, 33].

In the experimental setup of Watters et al. [16] and Yu et al. [15], the SHAPE reagent is added approximately 30 s after transcription stopped at the different transcript lengths. The probing reagent benzoyl cyanide (BzCN) itself can be considered very fast, as the reaction happens on the order of 250 ms (an order of magnitude slower than transcription, but negligibly fast compared to the 30 s delay) [34]. Thus, we start separate `DrTransformer` simulations for each transcript length and set the simulation time for the last nucleotide to 30 s. The simulated SHAPE profile is then composed from the end of simulations for each transcript length.

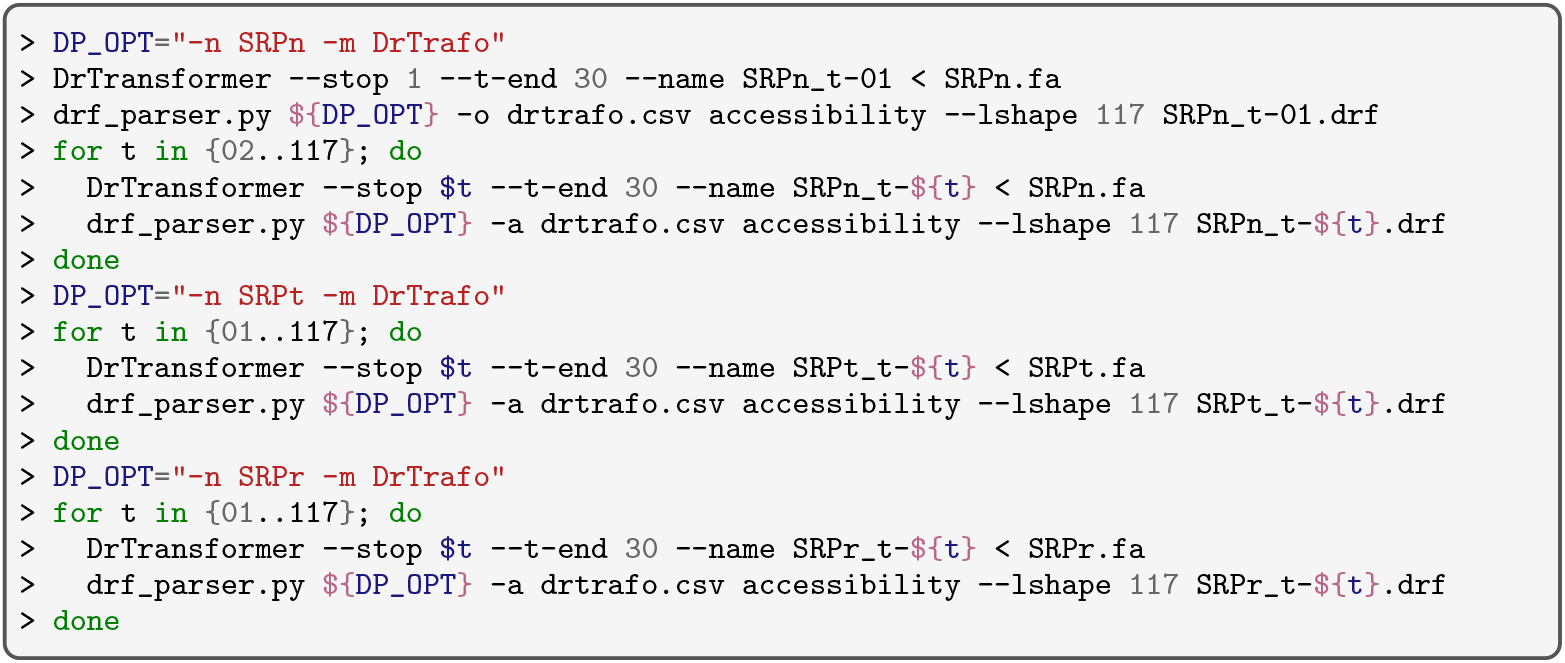

#### 3.9.2 Preparation of cotranscriptional SHAPE data

Cotranscriptional SHAPE reactivity data for SRPn, SRPt and SRPr is available at the RNA Mapping DataBase (https://rmdb.stanford.edu/) as entries `SRPECLI_BZCN`, `SRPU21C_BZCN`, and `SRP21CR_BZCN`, respectively [16, 15]. For convenience, we provide a copy of this data in the `SHAPE/` directory of our git repository. We further use the script `convert_rdat.py` to convert the `RDAT`-formatted files into the same `CSV` format as that from Sec. 3.7:

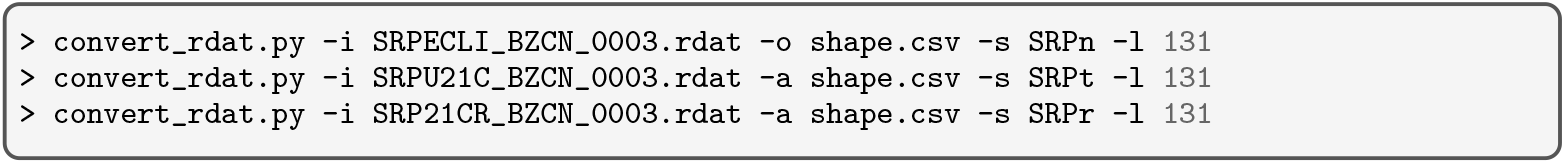

Note that here we use the experimental replicate 3, but we encourage the reader to repeat this analysis with the other provided files, or with the equilibrium refolding SHAPE experiments available for SRPn and SRPt. In contrast to data for the mutant SRP sequences the SRPn data does not cover the full length transcript when including the polymerase footprint. To compare SHAPE results that cotain a different number of transcripts, we provide the command line option `-l 131` that sets the expected transcript length to 131 nt (117 nt SRP plus 14 nt polymerase footprint) for each input file. Finally, all extracted data can be stored in the file `shape.csv`, where any missing data is indicated by `NA` values.

#### 3.9.3 Plotting and interpreting the results

The following commands utilize the `plot_accessibility.R` script to plot both an accessibility and a reactivity profile for the cotranscriptional SHAPE data and the corresponding `DrTransformer` simulations:

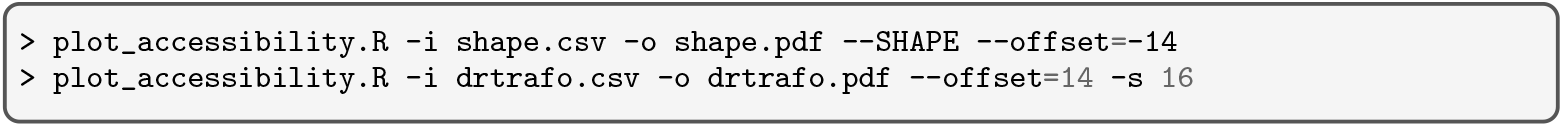

In the first call of the plotting script, we provide the `--SHAPE` option to indicate that the input data is composed of SHAPE reactivities. It is reasonable to assume that SHAPE reactivities *ρ* ≥1 originate mostly from unpaired positions. Our script by default already crops the reactivity data accordingly. Note, however, that in the publications of Watters et al. [16] and Yu et al. [15] a threshold of *ρ*≥ 4 has been applied, see also Note 7. The `--offset` option adjusts the labels on the y-axis of the plot. SHAPE data taken from Watters et al. [16] and Yu et al. [15] includes a 14 nt polymerase footprint which does not participate in cotranscriptional folding, hence the argument of −14. Simulated data on the other hand does not include a polymerase footprint, so we apply a positive offset of 14 nt to only adjust the y-axis labels for the sake of better comparison against the SHAPE data. The option `-s 16` starts the visualization at a transcript length of 16 nt. This option is not required for plotting the SHAPE data since no data for shorter sequences is available. Both resulting plots will be stored in the files `shape.pdf` and `drtrafo.pdf`, respectively.

The visual comparison of SHAPE-Seq reactivities against simulated accessibilities in Fig. 6 shows that experimental reactivities are generally lower. As expected, the last 14 nt of the SHAPE-Seq data are mostly inaccessible due to the polymerase footprint. Surprisingly, also the presumably unstructured regions close to the 3^*1*^ end of the nascent transcript show consistent patterns of (in)accessibility, (cf. “white triangle” in `DrTransformer` data between 40 and 60 nt transcript length). Also the experimental reactivity patterns that correspond to H1 formation already start before H1 formation is even possible, i.e. when the nascent RNA is only 16 nt long. These effects may be consequences of partial detachment of the polymerase from the transcripts prior probing, i.e. revealing the otherwise inaccessible footprint.

Helix H1 – between positions 3 and 25 – can be seen throughout all experiments as parallel lines with low reactivity, resp. accessibility. In many cases this includes the two additional base-pairs enclosed at the tip of H1 (see also Fig. 1B). At about 115 nt transcript length, both SRPn and SRPr show lower reactivities within the unpaired regions of H1 hairpin, presumably because many of the probed molecules form alternative structures instead, e.g. the functional H2-S1-S2-S3-S4 conformation.

The change in reactivities at about 80–85 nt transcript length in SRPt and SRPr can be related to the predictions with `DrTransformer`, where H2 is formed as a precursor towards the functional structure, see also Fig. 1F. Unfortunately, the final refold to the rod-like functional form of the SRP structure (H2-S1-S2-S3-S4) is not clearly visible in the SHAPE data. The accessibility patterns in the H1 region change for all three sequences when roughly 110 nucleotides are available for folding, but it is not visible by eye which structures are present in the ensemble. A more detailed analysis is beyond scope of this contribution, but necessary to minimize the danger of cherry-picking suitably patterns from reactivity profiles. E.g. reactivity profiles can also be interpreted by tracking relative changes of reactivity – so-called upswing/downswing events [35].

### 3.10 Tracing cotranscriptional motif formation

The final analysis method in this tutorial assumes that we know a set of particular motifs to search for, e.g. revealed by visual inspection as in Fig. 4, or also via experimental data as in Fig. 6. The formation of selected motifs in the ensemble can then be traced over time in the output of `DrKinfold` or `DrTransformer`, the results are shown in Fig. 7.

**Figure 7:**
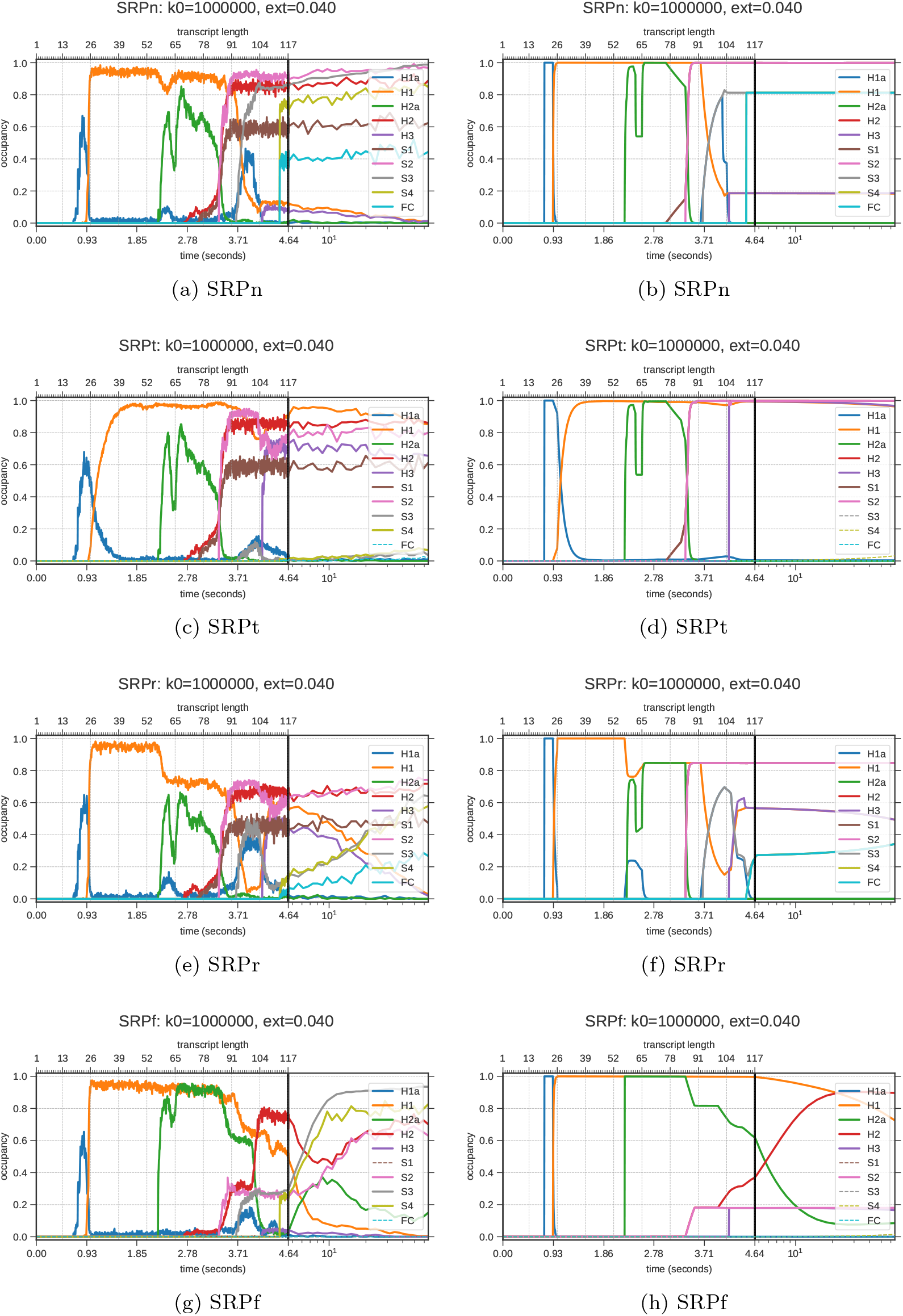
Occupancy of target motifs over time in `DrKinfold` and `DrTransformer` output. The vertical black line marks the transcription of the last nucleotide and also the transition from a linear time scale to a logarithmic time scale. The end point marks 60 seconds after the last nucleotide has been transcribed. All plots compare motifs described in Fig. 1. Here, FC is defined as the union of motifs H2, S1, S2, S3 and S4.

Generally we prioritized motifs that are interesting in all of the different SRP variants for visualization in Fig. 7. Specifically, we trace motifs enclosing hairpins: H1a, H1, H2a, H2, H3, and the stems required for the rod-like conformation: S1, S2, S3, and S4 (see Fig. 1). We also show the functional conformation, FC, which we define as the union of motifs H2, S1, S2, S3, and S4. A summary of investigated motifs in dot-bracket notation can be found in the file `sequences/SRP.mot`. Note that this file contains additional motifs not shown here, such as H1b, H1c, H3a, PK and PK2.

Among the motifs *not* shown in Fig. 7 is H1c, a motif suggested by Fukuda et al. [14] to compete with H1, as well as the motifs PK and PK2 to find evidence for a pseudoknot interaction suggested by Yu et al. [15]. Both motifs PK and PK2 identify structures with six simultaneously unpaired bases. If those bases are unpaired at the same time, that would allow for a three-basepair pseudoknotted toehold interaction. However, the results are not shown, as we found no compelling evidence for this scenario.

#### 3.10.1 Generating motif plots

We repeat the `DrTransformer` and `DrKinfold` analysis from Sec. 3.6.1 with the option `--t-end 60` to follow the formation of motifs for a minute after transcription. The calls are shown below:

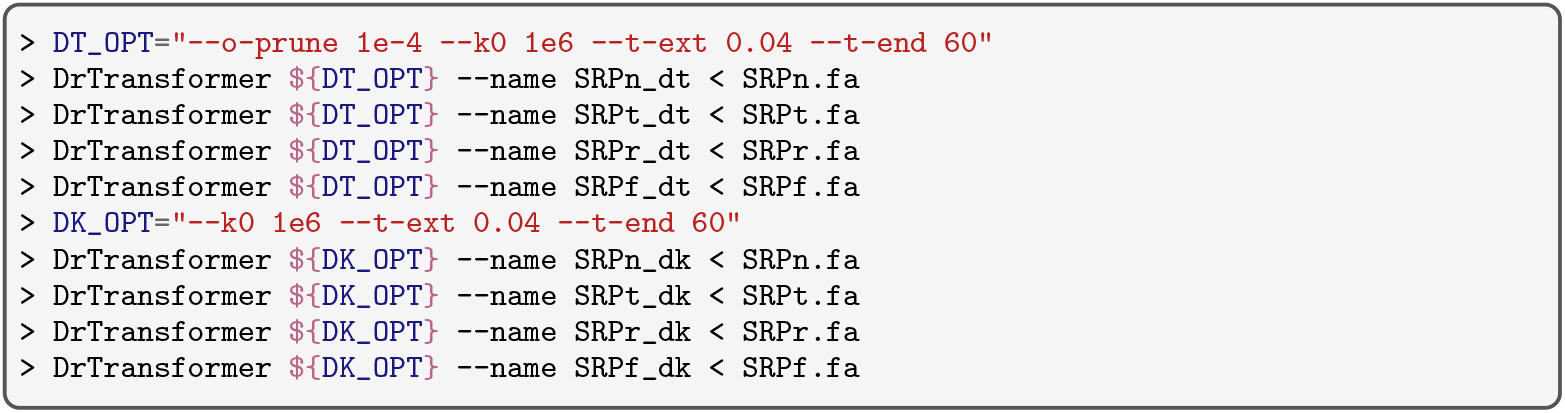

The plots to trace motifs can be generated using the `DrPlotter` program that is installed together with `DrTransformer`. We need to specify the file that contains motif specifications, as well as a list of motifs that defines which (and in which order) motifs should be plotted.

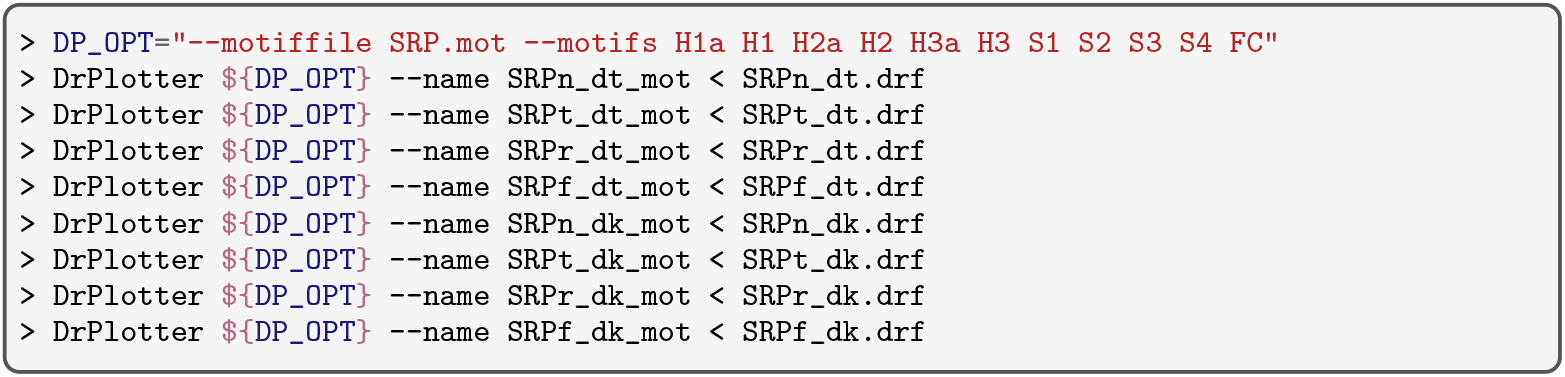

#### 3.10.2 Motif plot interpretation

The plots in Fig. 7 compare `DrKinfold` and `DrTransformer` predictions. As usual `DrKinfold` is the more detailed model, but `DrTransformer` output can also be insightful, as it ignores base-pair level variations.

For the native SRP molecule, we see that H1a and H1 are in competition. H1a forms first as it is the first cotranscriptionally available motif, but it is quickly replaced by H1. However, around 95–100 nt length when the S3 stem forms, H1a becomes the dominant structure again. Further visual inspection (with `DrForna`, not shown) reveals a causal relationship, as the H1a hairpin is compatible with S3 formation, and the newly transcribed part of the RNA can invade the stem of motif H1a to gradually extend S3 – a relatively fast process also known as three-way branch migration.

Furthermore, H2a precedes the formation of H2, and less than 1% of structures form H3 at any point in time. The motifs H2, S2, S3, and S4 are approaching more than 80% in the simulated ensemble after transcription, only S1 is less occupied at 60%. The union of all constraints, FC, is at about 40% occupancy after transcription. The known functional structure is less dominant, as it is unlikely that all enforced base-pairs are closed at the same time. In the `DrTransformer` output we see that essentially all structures form S1 and S2, but only 80% form S3 and S4, as the remaining 20% of the ensemble is trapped with H1 and H3. Accordingly, FC reaches 80% (the line overlaps with formation of S4).

Not shown in Fig. 7 is the motif H1b, which is only present during the transitions from H1a to H1 and from H1 to H1a. H1c is never formed, and there is no evidence for a pseudoknotted reconfiguration, as the bases that would initiate the interaction (PK and PK2) are almost never accessible at the same time.

The trapped SRPt molecule forms mostly H2-S1-S2 in combination with H1 and H3, and cannot rearrange into the functional FC form. The rescue mutant SRPr shows an interesting competition between H1 and H1a, but eventually all helices of FC do form at the end of transcription in 10−− 20% of structures. The H1/H3 structure is still at about 50% at the end of transcription, with occupancy rapidly declining within 60 seconds.

The SRPf sequence forms H2, S2, S3 and S4 at the end of transcription, but the mutations inhibit the formation of S1. Thus, it appears as if FC did not form, even though most of its structure is still intact. Interestingly, `Kinfold` predicts a second round of H2a formation after transcription, when H1 is dissolved. Note that `DrTransformer` predictions are not simply less noisy predictions compared to `Kinfold`. Here, H1 remains stable until the end of transcription, and only the S2 stem forms at 20%. All other motifs for FC are not observed on the investigated time scale. The formation of H3a is not shown, but it is predicted by both `Kinfold` (60%) and `DrTransformer` (80%) at around 90 nt length.

### 3.11 Concluding remarks

We have shown how to use computational methods to find structural motifs that are present at specific points in time during cotranscriptional folding. Of course, the underlying thermodynamic nearest neighbor model makes certain simplifying assumptions about which structures are allowed and what their free energy is. Moreover, errors in free energies are exponentially amplified when calculating transition rates between structures (see Eq. 1). However, we showed that computational models can be useful to help with the interpretation of experimental data. We present results with *k*_0_ = 10^6^ s^−1^ in this tutorial, even though related literature used *k*_0_ = 10^5^ s^−1^ for `Kinfold` related kinetic models [36, 37]. The main reason for choosing *k*_0_ = 10^6^ s^−1^ is the correspondence of simulations with experimental data, as we were not able to observe the same nice fits with *k*_0_ = 10^5^ s^−1^. However, this parameter also makes sense, as it is closer to the fastest reaction rates known for RNA folding.

The software discussed here is only a part of generally available cotranscriptional folding software. Among other cotranscriptional folding simulators are `Kinefold` [38], `RNAkinetics` [39], `Kinwalker` [40], `BarMap` [41], and `CoStochFold` [42]. The `drconverters` repository provides a script to convert results from `Kinefold` to the `.drf` file format. `Kinefold` is especially interesting as it allows for pseudoknotted conformations, but, unfortunately, uses outdated nearest neighbor parameters.

We modeled the cotranscriptional process as continuous process that attaches nucleotides at equally spaced time intervals. However, nucleotide attachment is a stochastic process and there are known pause site motifs in the SRP sequence. In principle, such pause sites could have been considered for cotranscriptional simulations (e.g. see `DrTransformer` command line parameters). For the SRP pause sites suggested in literature we have not found a large influence in structure formation. This is because the pause sites are at nucleotides where the ensemble is close to the equilibrium distribution, such that there are no notable impacts on downstream folding events.

While we only compared against cotranscriptional SHAPE data here, it is relatively straight forward to plot the length of the exterior loop and compare those results with the optical tweezer results from [14]. However, an interpretation of those results seemed even more difficult without direct correspondence with experimenters and thus was beyond scope for this tutorial. We hope that this tutorial provides a foundation to investigate many more related systems and questions.

## 4 Notes

1. A PC with `Windows` 10 or above may work in combination with the Windows subsystem for Linux (WSL). Native Windows support without WSL is not available for the tasks presented within this chapter, since the required `ViennaRNA Package` Python bindings are at this time not available for Windows.
2. Instead of installing the scripts globally, they can be installed on a per-user base by providing `pip` with the flag `--user`:

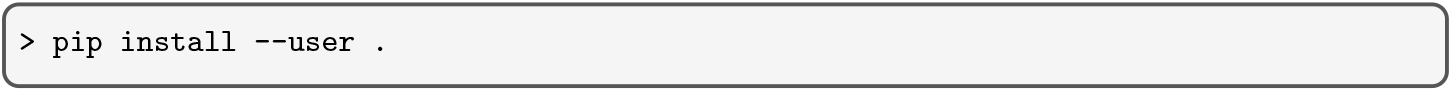 If installation of scripts is not desirable, then all executions must be adapted to include location of scripts on disk. For instance the command

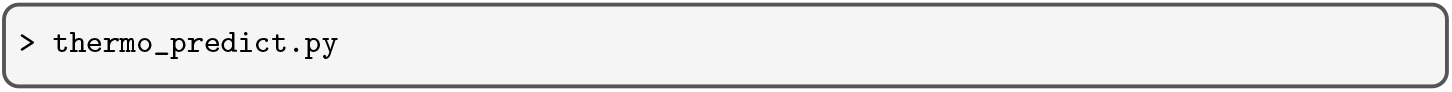 becomes

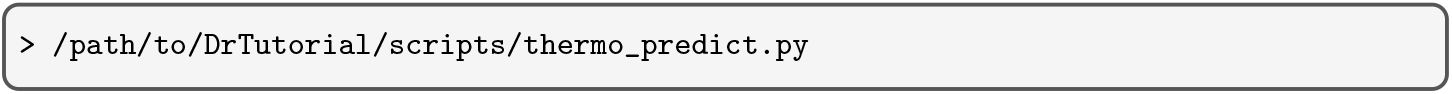 where `/path/to/DrTutorial` is the absolute path to the DrTutorial directory. Alternatively, users can set the `PATH` environment variable such that it includes the absolute path to the `scripts` directory, e.g.

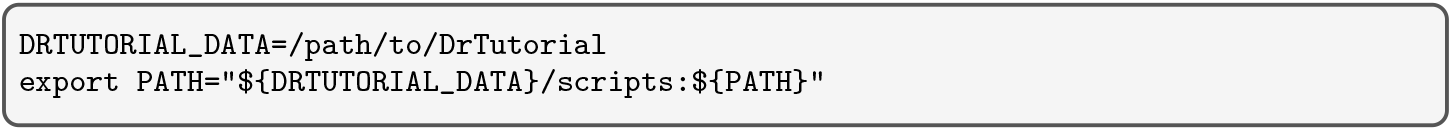 To make the environment variable permanently available throughout different `Unix` terminal sessions, you may store the above lines in the appropriate config file of your terminal emulator, for instance the `bash` terminal emulator uses the `∼/.bashrc` file.
3. Beware of the fact that most RNA secondary structure prediction programs like `RNAfold` only backtrack *a single* MFE structure. However, some RNA sequences can form multiple distinct secondary structures with minimum free energy. To enumerate *all* MFE structures with `RNAsubopt` use:

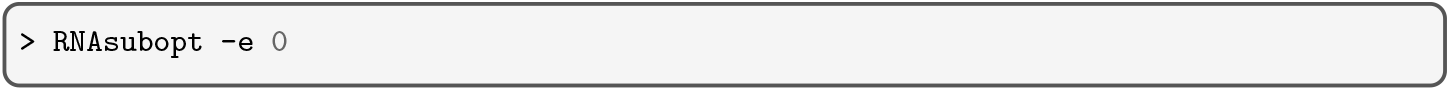 where the `-e 0` option tells `RNAsubopt` to print all structures within a 0 *kcal · mol*^−1^ energy range above the MFE.
4. By default, the `-p/--stochBT` option of `RNAsubopt` yields a list of secondary structures *s* from the equilibrium distribution, randomly drawn according to their probability *p*(*s*). The free energy of these structures is not part of the output due to implementation details. However, the `RNAsubopt` program comes with a second, alternative, option `--stochBT_en` that can be used as a substitute. As a result, a program call like:

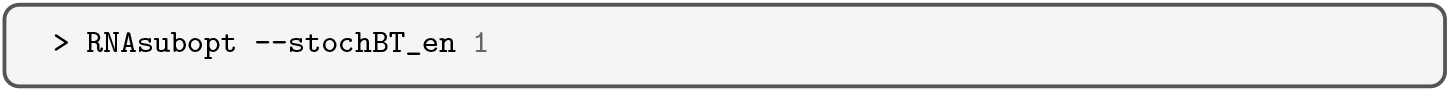 yields a single secondary structure followed by its free energy *E*(*s*) in units of kcal *·* mol^-1^ and the corresponding equilibrium probability 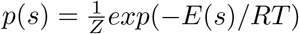 with temperature *T*, gas constant *R* and partition function *Z* = *s exp*(−*E*(*s*)*/RT*).
5. Since any nucleotide *i* may pair with any other nucleotide *j*, the number of possible base pairs (*i, j*) grows quadratic in the length of the sequence. Consequently, a visualization of the pairing probabilities during transcription or the change thereof would require a 4− dimensional plot: two dimensions to address the actual base pair, one for the length of the nascent transcript, and one for the actual pairing probability.
6. Base pair probabilities *p*_*ij*_ can be readily computed using a command line call like:

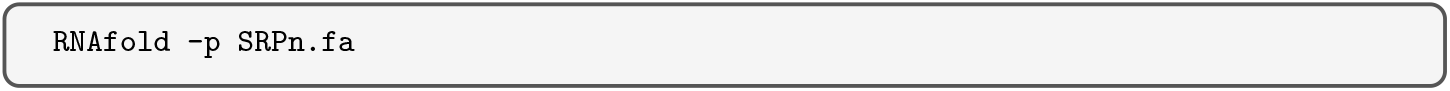 This will create a *dot-plot* PostScript output file `SRPn_dp.ps` that provides a visualization of the individual pairing probabilities and stores them in a machine readable form for further processing. For each base pair (*i, j*) and corresponding probability *p*_*ij*_ the PostScript file contains lines of the form

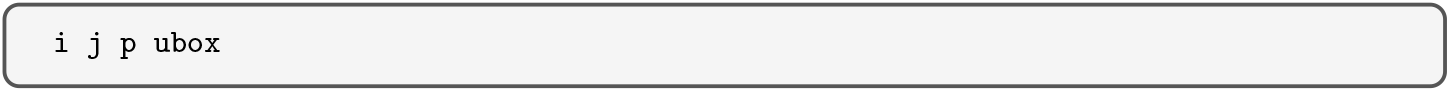 where *i* and *j* are integer numbers denoting the two positions that pair (1-based), and *p* is a floating point number showing the square-root 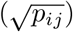 of the corresponding pairing probability *p*_*ij*_.
7. For the cotranscriptional SHAPE reactivity profiles shown in Fig. 6 we applied a threshold of *ρ* ≤1. This cropping of the data together with the limits of the color scale from 0–1 may conceal the full picture of potential conformations probed by the SHAPE reagent. To change the cutoff value, the `plot_accessibility.R` script takes an optional argument `--SHAPErange r` where *r* is the largest value of *ρ* that should be considered. For instance, to replicate the data shown in Watters et al. [16] and Yu et al. [15] one may use a range of 0≤ *ρ*≤ 4 by adding the parameter `--SHAPErange 4` upon creating the plot.

## Footnotes

1 https://viennarna.github.io/drforna/

